# The biochemical function of bivalent aptamer assemblies against B-cell markers CD19 and CD20

**DOI:** 10.1101/2025.01.26.634939

**Authors:** Yilam Ng Cen, Nicole B. Williams, Nicolás Di Siervi, Lexi Chen, Pramodh Seneviratne, Geri Kreitzer, Leandro Cerchietti, Prabodhika Mallikaratchy

**Affiliations:** Ph.D. Program in Chemistry and Biochemistry, CUNY Graduate Center, The City University of New York, 365 Fifth Avenue, New York, NY 10016; Ph.D. Program in Biology, CUNY Graduate Center, The City University of New York, 365 Fifth Avenue, New York, NY 10016; Department of Molecular, Cellular, and Biomedical Sciences, CUNY School of Medicine, City University of New York, New York, NY10031; Department of Medicine, Division of Hematology and Oncology, Weill Cornell Medical College, 1300 York Ave., room C-620-B, New York, NY10065

## Abstract

Aptamers are synthetic oligonucleotides that bind to their specific receptors with high specificity, offering immense potential for the development of molecular tools. Using recently introduced Ligand-Guided Selection (LIGS), a variant of the Systematic Evolution of Ligands by Exponential Enrichment (SELEX) we previously identified anti-CD19 and anti-CD20 aptamers with specificity and affinity. Given their high expression levels, B-cell markers CD19 and CD20 are widely utilized in the diagnosis of B-cell-related malignancies and autoimmune diseases. Here, we report the design and functional characterization of bivalent aptamer assemblies targeting CD19 and CD20 expressed in B-cell lymphomas. Using a strategic approach, we synthesized dimeric constructs of these aptamers with polyethylene glycol (PEG) linkers of varying lengths to tether the two aptamer units. The bivalent aptamers demonstrated enhanced binding affinity and specificity, with an optimal linker length of ∼3.96 nm. Functional studies revealed that dimeric CD19 aptamers selectively internalized in CD21-negative B-cells, while CD20 aptamers exhibited improved antigen binding without triggering calcium release. These findings highlight the potential of bivalent aptamers in engineering cost-effective, stable, and precise therapeutic agents for B-cell-related malignancies, such as diffuse large B-cell lymphoma (DLBCL). This work advances the development of aptamer-based synthetic therapeutics with promising clinical applications.

## Introduction

B-cells are central to the adaptive immune system and are implicated in many diseases, including B-cell malignances, such as non-Hodgkin’s Lymphoma (NHL) and Hodgkin’s Lymphoma (HL), as well as autoimmune diseases, such as systemic lupus erythematosus (SLE), multiple sclerosis (MS), and rheumatoid arthritis (RA).^1, 2^ As a result, B-cells markers have been recognized as critical antigens in the development of therapeutic strategies.^2^ Two widely targeted B-cell-specific antigens are CD19 and CD20. CD20, or cluster of differentiation 20, is a protein found on the surface of B-cells; it is involved in B-cell development, differentiation and B-cell receptor (BCR) signaling, and it is found in B-cell lymphomas and leukemias.^3^ A monoclonal antibody known as Rituximab has been extensively used to treat B-cell associated malignancies.^3^ Importantly, the use of alternative B-cell targeting antibodies that recognize different antigens, such as CD19, has shown success in overcoming Rituximab resistance in CD20-negative B cells.^4^ CD19 is, therefore, a valuable target in treating B-cell-related malignancies and other diseases. Moreover, the restricted expression of CD19 and its critical role in B-cell function also make it a valuable target for precision therapies. Additionally, therapeutic strategies, such as chimeric antigen receptor (CAR) T cells and bispecific antibodies targeting CD19, have shown remarkable success in treating B-cell cancers by promoting immune-mediated tumor cell destruction.^5^ While monoclonal antibody (mAb) strategies have been successful in the development of targeted therapies, challenges associated with mAb production, as well as low temperature and pH stability, have limited their wider applicability.^6–8^

Aptamers are comprised of short single-stranded DNA or RNA that folds into a three-dimensional structure which facilitates specific target binding. Several characteristics unique to aptamers make them superior to antibodies, e.g., small size, flexibility, and ease of chemical modification.^9–11^ Aptamers have recently shown therapeutic success. For example, pegaptanib, an aptamer against vascular endothelial growth factor for treating age-related macular degeneration (AMD), was approved by the U.S. FDA in 2004.^12^ A second aptamer against AMD was approved by the FDA in 2023, and several promising aptamer candidates against various disease targets are currently under clinical trials, demonstrating the potential of aptamers as therapeutics.^9, 12^ Aptamers could also be utilized to develop synthetic antibodies for B-cell-related diseases. Accordingly, given the significance of CD19 and CD20 in disease-related B-cells, we recently utilized two monoclonal antibodies and identified multiple aptamer candidates against CD20 and CD19 with high specificity. These aptamers were identified using a novel method called multi-target Ligand Guided Selection (LIGS) using a cell-SELEX (Systematic Evolution of Ligands by Exponential enrichment) library enriched against Toledo, a B lymphocyte cell line.^13, 14^

It has been established that bispecific antibodies show promise in the treatment of several malignancies, such as non-Hodgkin’s Lymphoma and multiple myeloma.^15^ In light of this, we herein report the development of dimeric aptamers against CD19 and CD20 with the potential to be utilized in engineering bispecific aptamers. More specifically, aptamers represent synthetic, single-stranded DNA molecules that act like a antibody by specifically binding to a target molecule with high affinity, thus opening the door to generate stable, cost-effective therapeutics.^16^ After carrying out the structure-activity relationship (SAR) of aptamers, we previously reported that truncated aptamers could be reengineered into dimeric versions to increase their affinity and functionality.^17–20^ We further showed that truncation of aptamer sequences would not impair their ability to fold into tertiary structures or function normally.^17–20^ Therefore, based on our prior studies, we herein truncated anti-CD20 and anti-CD19 aptamers into shorter variants by removing nonessential regions, while preserving both structural integrity and binding affinity. Truncated variants with the highest affinity and specificity were then used to design dimeric aptamers composed of two different aptamers. Finally, the dimeric variant having the most optimized linker was investigated for internalization and specific function against CD19 and CD 20. We found that the dimeric aptamers against CD19 slowly internalize in CD21-negative CD19-positive lymphoma cells, cancerous B-lymphocytes that express the CD19 marker, but lacking the CD21 marker, suggesting their applicability in designing synthetic bivalent therapeutics against subset of human lymphoma. Meanwhile, dimeric CD20 aptamer showed improved antigen binding, but it did not induce calcium release.

## Results

### Truncation and binding affinity analysis of anti-CD19 and anti-CD20 aptamers

We reported two aptamers against CD19, namely WB15-CD19 and WB17-CD19. Two mutations distinguish the two aptamers, but both share similar sequence identities. We previously designed anti-CD3 aptamers and observed a similar trend whereby a family of aptamers originating from a single sequence with mutations could be identified using LIGS. Based on this observation, we focused on truncating both aptamers to generate shorter variants. As previously noted, we and others have utilized systematic truncation strategies to produce shorter variants of aptamers selected against targets on whole cells. Therefore, following a similar approach, we first truncated 8 bases from the 3’ end and 4 bases from the 5’ end, removing a total of 12 bases to generate the WB15.CD19.1 variant, which consists of 65 bases (**Table 1; Fig. S1A-C; Fig. S2A-B**). Assuming that the aptamer’s functional region forms a hairpin, we further removed 22 bases to generate WB15.CD19.2. Both variants were assessed for their affinity constants. Interestingly, the WB15.CD19.1 variant exhibited a slightly decreased affinity, while WB15.CD19.2 completely lost its binding capability towards CD19 expressed on Raji cells. This observation suggests that the aptamer binding mechanism might follow an “induced fit” model, rather than a “fold-and-bind” model, thus restricting the ability to further truncate these aptamers to generate even shorter variants (**Table 1; Fig. S1A-C; Fig. S2A-B**). Using a similar approach, we truncated WB17-CD19 by removing bases from the 3’ and 5’ ends to generate a 65-mer termed WB17.CD19.1. However, a trend similar to that observed for the WB15.CD19 emerged, and the WB17.CD19.2 variant also lost its binding ability, indicating that neither aptamer could be truncated further without disrupting functional folding (**Table 1; Fig. S1E-G; Fig. S2C-D**).

**Table 1:**
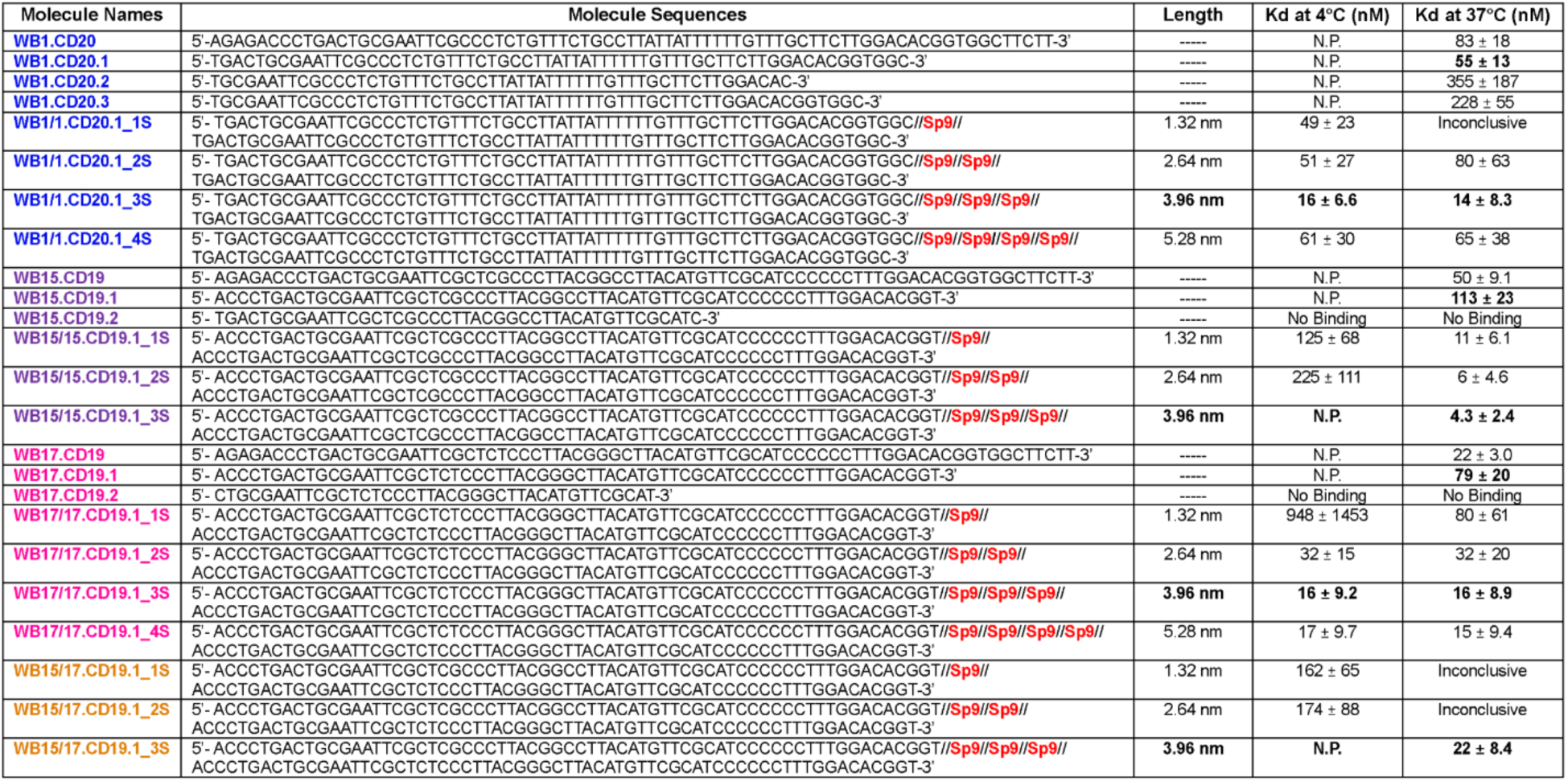
Apparent binding Affinities (apparent Kd) of Monovalent and Dimeric CD19 and CD20 Aptamers. This table summarizes the apparent binding affinities (Kd values) of monovalent aptamers (WB1.CD20, WB15.CD19, WB17.CD19) and their truncated variants, as well as dimeric aptamers constructed using the best-performing monovalent truncations tethered by spacer 9 PEG linkers. apparent binding affinities were determined using Raji cells (Burkitt’s lymphoma) at concentrations ranging from 500 nM to 1.95 nM for monovalent aptamers and 250 nM to 0.1 nM for dimeric aptamers. Data are presented as mean ± SD from three independent experiments. Dimeric aptamers demonstrated significantly improved apparent binding affinities compared to their monovalent counterparts with optimal affinity observed for WB15/15.CD19.1_3S, WB17/17.CD19.1_3S, WB15/17.CD19.1_3S, and WB1/1.CD20.1_3S variants.

Next, we focused on the previously reported CD20 aptamer, WB1.CD20, to generate shorter variants. Using an approach similar to that described above, we first removed 12 bases—4 bases from the 3’ end and 8 bases from the 5’ end—to generate WB1.CD20.1, which consists of 65 bases (**Table 1; Fig. S3A-D; Fig. S4A-D**). We further truncated 9 more bases to form WB1.CD20.2 and WB1.CD20.3. Both shorter variants lost their affinity towards CD20 (**Table 1; Fig. S3A-D; Fig. S4A-D**). However, the WB1.CD20.1 variant did show improved binding affinity towards CD20 expressed in Raji cells. Attempts to truncate WB2.CD20 to generate shorter variants were unsuccessful (**Fig. S4E-F**).

### Specificity of truncated anti-CD19 and anti-CD20 aptamer variants

The truncated variants were then assessed for their specificity using CD19/CD20-positive and CD19/CD20-negative cells, including knockout CD19 and CD20 Burkitt’s lymphoma CA46 cells. Since only the first truncation showed considerable binding affinity, we focused on assessing the specificity of WB1.CD20.1 aptamer against CD20-positive cell lines, including the Burkitt’s lymphoma cell lines Raji, BJAB, SKLY-16, and the Non-Hodgkin’s Lymphoma cell line Toledo. Negative control cells included T-cell leukemia Jurkat.E6 and CD20-knockout (CD20KO) CA46 cells (Burkitt’s lymphoma). Based on the binding patterns towards positive and negative cell lines, the WB1.CD20.1 variant did not lose its binding specificity towards CD20 on the surface of B-cell lines (**Fig. S5A-G**).

Then, we utilized the same cell lines to assess binding specificity of CD19 variants WB15.CD19.1 and WB17.CD19.1. While we observed slightly decreased binding affinity of truncated CD19 variants compared to their parent aptamers, the specificity of the aptamers towards CD19 was unaffected by truncation, as demonstrated by specific binding of both variants towards the CD19-positive cell lines Raji, Toledo, BJAB, and SKLY-16, but not towards CD19KO CA46 cells or CD19-negative Jurkat E6.1 cells (**Fig. S5H-N and Fig. S5O-U**). The observed high specificity of truncated WB1.CD20.1, WB15.CD19.1, and WB17.CD19.1 suggests that truncation had no effect on the specificity towards their respective targets. Finally, using confocal microscopy, we next assessed whether truncated CD19 aptamer WB17.CD19.1 and truncated CD20 aptamer WB1.CD20.1 would colocalize with their respective monoclonal antibodies. As expected, both aptamers did colocalize with their respective antibodies, further demonstrating target specificity (**Fig. S6**).

### Dimerization of anti-CD19 and anti-CD20 aptamers

Dimerization is a process whereby two similar or identical molecules are tethered by bonds to form a single polymer called a homodimer. Since the apparent affinities of monomeric aptamers are insufficient for functional cellular assays, we dimerized both anti-CD20 and anti-CD19 aptamers to enhance their apparent affinity. The generation of dimeric aptamers to improve affinity and functionality is well established. Among the various available linkers, we used Polyethylene Glycol (PEG) linkers to tether both aptamers. Monomeric aptamers typically contain 35 to 45 bases, and longer PEG linkers are generally more suitable for dimerization within this size range. However, since both anti-CD19 and anti-CD20 aptamers are 65-mer, we employed spacer 9 to optimize linker length between the two aptamers. Four dimeric variants of WB1.CD20.1 were designed with linker lengths ranging from one spacer (∼1.32 nm) to four spacers (∼5.28 nm) (**Table 1; Fig. S3E**). The first dimeric variant, WB1.CD20.1_1S with a single spacer showed improved binding affinity (dissociation constant, Kd = 49 ± 23 nM) compared to the monomeric variants at 4°C (**Table 1; Fig. S3E; Fig. S7A**). The second variant, WB1.CD20.1_2S, with two spacers exhibited similar binding affinity (Kd = 51± 27 nM) (**Table 1; Fig. S7B**). Interestingly, the best binding affinity was observed with WB1.CD20.1_3S, containing three spacers (Kd = 16 ± 6.6 nM) (**Table 1; Fig. S7C**). In contrast, the fourth variant, WB1.CD20.1_4S with four spacers demonstrated reduced binding affinity (Kd = 61 ± 30 nM) (**Table 1; Fig. S7D**), suggesting that the optimal spacer length is ∼3.96 nm, corresponding to three spacers. When apparent binding affinities were assessed at physiological temperatures, all variants showed reduced affinity, indicating decreased stability of aptamers at higher temperatures (**Table 1; Fig. S7E-H**). For example, WB1.CD20.1_1S exhibited no binding at 37°C (**Fig. S7E**). However, binding gradually improved with longer spacer lengths, as observed with WB1.CD20.1_2S, which showed moderate binding at 37°C (Kd = 80 ± 63 nM) (**Fig. S7F**). WB1.CD20.1_3S maintained strong binding at 37°C (Kd = 14 ± 8.3 nM) (**Fig. S7G**), confirming its superior stability and performance across temperatures. Finally, WB1.CD20.1_4S displayed weaker affinity at 37°C (Kd = 65 ± 38 nM) (**Fig. S7H**).

Next, we used two truncated aptamer variants, WB15.CD19.1 and WB17.CD19.1, against CD19 and synthesized three sets of dimeric aptamers. First, we designed two homodimeric aptamers, each consisting of either two WB15.CD19.1 or two WB17.CD19.1 aptamers. Following a linker optimization approach similar to that used for anti-CD20 aptamers, we optimized the linker length by varying the number of PEG spacer units between the aptamers (**Table 1; Fig. S1D and S1H; Fig. S8**). Affinity of the homodimeric aptamers WB17/17.CD19.1_1S and WB15/15.CD19.1_1S tethered with a single spacer showed apparent Kd values of 948 ± 1453 nM and 125 ± 68 nM, respectively (**Table 1; Fig. S8A and S8I**). At physiological temperatures, the apparent Kd for both dimeric aptamers improved to 80 ± 61 nM and 11 ± 6.1 nM, respectively (**Table 1; Fig. S8E and S8K**). The apparent affinities for both dimers at 4°C and 37°C were further improved with the use of two spacers (T**able 1****; Fig. S8B/S8F and S8J/S8L**). The optimal configuration with the highest apparent affinity was achieved by using three spacers, resulting in a Kd of 16 ± 8.9 nM for WB17/17.CD19.1_3S and 4.3 ± 2.4 nM for WB15/15.CD19.1_3S (**Table 1; Fig. S8G and S8M**). Unlike the observation with anti-CD20 dimeric aptamers, the use of four spacers 9 did not lead to weaker binding. Instead, it maintained a similar binding affinity (Kd = 17 ± 9.7 nM) (**Table 1; Fig. S8D**) while plateauing at the three spacers 9, with no significant improvement observed with the extended linker of a fourth spacer 9 (**Table 1; Fig. S8D/S8H**). We also explored the possibility of tethering WB15.CD19.1 and WB17.CD19.1 aptamers together to create heterodimeric aptamers. Using the same linker optimization strategy, we synthesized three variants with one, two, and three spacers (**Table 1; Fig. S8N-R**). However, the apparent binding affinities of the heterodimers were lower than those of the homodimers.

### Specificity of dimerized anti-CD19 and anti-CD20 aptamers

Following the optimization of linker lengths for dimeric anti-CD19 and anti-CD20 aptamers, we next evaluated their specificities. We used five lymphoma cell lines known to express CD19 and CD20 as positive controls: Burkitt’s lymphoma cell lines Raji, BJAB, Ramos, SKLY-16, and Toledo cells. As negative controls, we included Jurkat.E6 cells and CD19/CD20 knockout cells. Specificity was assessed at 37°C. First, we analyzed the specificity of the WB1/1.CD20.1_3S dimeric aptamer and observed that it was retained, despite dimerization (**Fig. 1A1-A7; B**). Similarly, we evaluated the specificity of three anti-CD19 constructs, each with three spacer units. All three constructs showed specific binding to all five cell lines expressing CD19, and no binding was observed to the CD19-negative cells (**Fig. 1C1-C7; D; E1-E7; F; G1-G7; H**). We further assessed the specificity of two promising dimeric aptamers, WB1/1.CD20.1_3S and WB17/17.CD19.3S, towards their respective antigens by performing colocalization studies with anti-CD20 and anti-CD19 monoclonal antibodies (mAbs). Confocal microscopy analysis of WB1/1.CD20.1_3S and anti-CD20 antibody on the Raji cell membrane showed that anti-CD20 aptamer and anti-CD20 antibody had colocalized on B-cells (**Fig. 1I-L**). Similarly, colocalization of WB17/17.CD19.1_3S with anti-CD19 antibody on Raji cells was observed, further confirming that the specificity of the truncated, dimerized aptamers had been retained (**Fig. 1M-P**).

**Figure 1:**
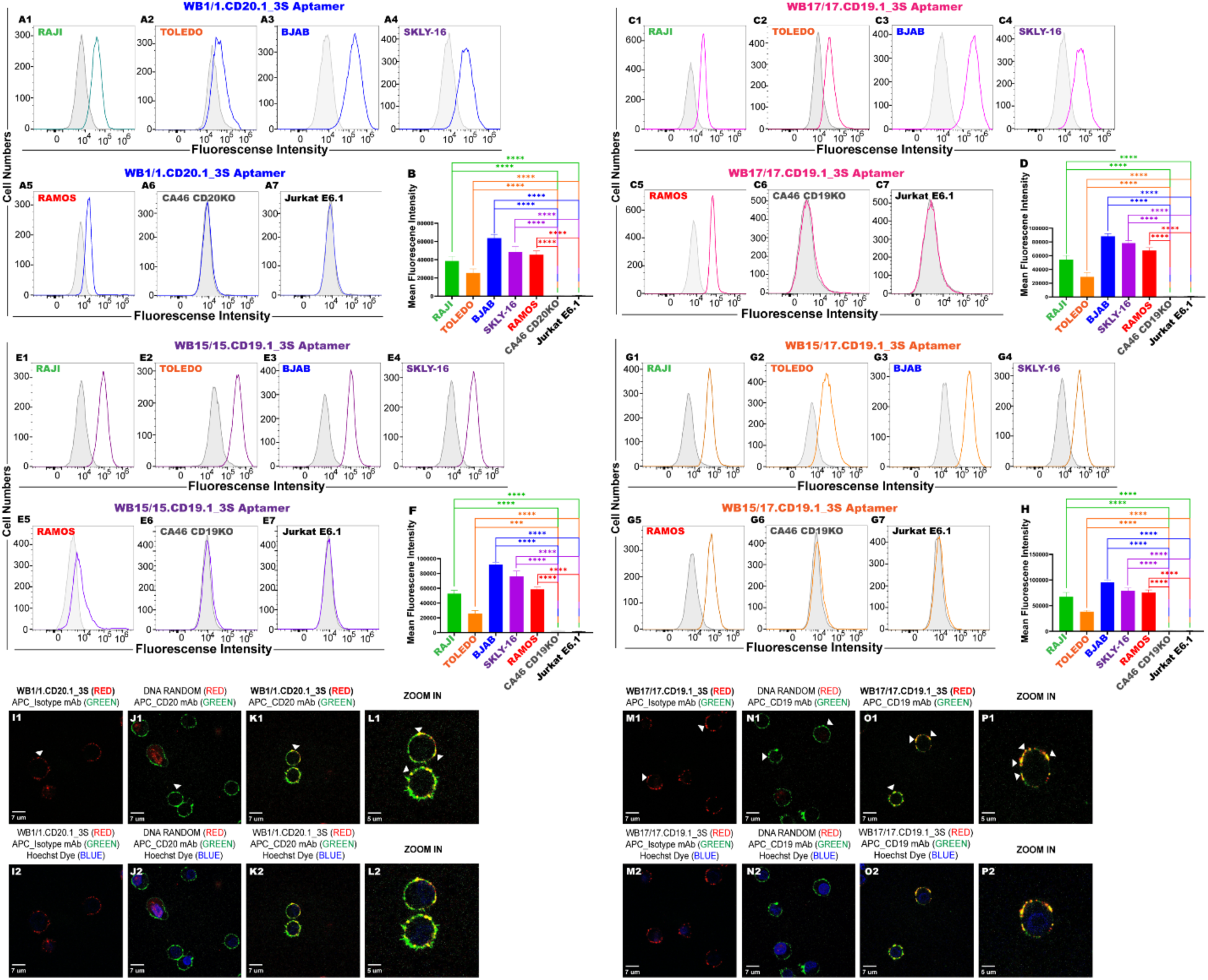
Specificity and Binding of Dimeric Aptamers Targeting CD19 and CD20 on B-cell lines. (A) Representative flow cytometric histograms showing the specificity of dimeric CD20 aptamer WB1/1.CD20.1_3S across CD20-positive cell lines (Raji, Toledo, BJAB, SKLY-16, and Ramos). (B) Bar graph quantifying the mean fluorescence intensities of WB1/1.CD20.1_3S binding, highlighting significant specificity (*p < 0.0001). (C; E) Fluorescence histograms for CD19 homodimeric aptamer WB17/17.CD19.1_3S and WB15/15.CD19.1_3S, respectively, demonstrating selective binding to CD19-positive cell lines (Raji, Toledo, BJAB, SKLY-16, and Ramos). (D; F) Quantification of homodimeric aptamer WB17/17.CD19.1_3S and WB15/15.CD19.1_3S, respectively, binding specificity using mean fluorescence intensity. (G) Fluorescence histograms for CD19 heterodimeric aptamer WB15/17.CD19.1_3S, demonstrating selective binding to CD19-positive cell lines (Raji, Toledo, BJAB, SKLY-16, and Ramos). (H) Quantification of heterodimeric aptamer WB15/17.CD19.1_3S binding specificity using mean fluorescence intensity. (I-L) Confocal microscopy images showing colocalization of bivalent WB1/1.CD20.1_3S (L1-L2: Cy3, RED) with a CD20-specific antibody (J1-J2: APC, GREEN) on Raji cells. DNA random aptamer controls and isotype antibody controls confirmed specificity. The aptamer binds to the cell surface membrane, as shown in zoomed-in views (L1-L2). Panel M-P: Confocal microscopy images of bivalent WB17/17.CD19.1_3S (M1-M2: Cy3, RED) colocalizing with a CD19-specific antibody (N1-N2: APC, GREEN) on Raji cells. The aptamer demonstrates specificity and surface binding with no significant off-target interactions. Zoomed-in views (P1-P2) confirmed aptamer binding to the cell membrane. Scale bars = 7 and 5 μm. Mean fluorescence intensity was calculated using the formula: Mean fluorescence intensity=Aptamer Mean Fluorescence − Random DNA Mean Fluorescence. The aptamer and random mean fluorescence values correspond to the mean fluorescence observed in their respective histograms. Bar graphs represent mean ± standard deviation from three independent experiments with statistical significance indicated (****p < 0.0001). Data represents mean ± standard deviation from three independent experiments.

### Internalization kinetics of dimeric CD19 aptamers

CD19 is a rapidly internalizing antigen.^21^ Given that dimeric CD19 aptamers could serve as highly effective synthetic drug delivery agents for CD19-positive human lymphoma, we next evaluated whether the aptamers could internalize into CD19-positive B-cells. To explore aptamer internalization, Xiao et al. used partial digestion of the extracellular portion of the receptor protein with trypsin.^22^ However, the use of trypsin is limited by its potential to act as a B-cell stimulator.^23^ Proteinase K is a cleaving reagent used to remove membrane-bound protein receptors.^24^ Therefore, as an alternative approach to assess aptamer internalization in B-cells, we used Proteinase K to partially digest the extracellular membrane portion of cells after incubation, followed by subsequent detection of internalized aptamers using flow cytometry. This method has been successfully used in aptamer binding analysis, but it has not been employed to explore aptamer internalization into cells.^25^ Since cell lines vary in the concentration of antigens on their surface, we first optimized Proteinase K cleavage of aptamers on the cell surface. CD19 antibody was used as a positive control to study the effect of Proteinase K treatment on CD19 antibody by fluorescence signal. The fluorescence intensity was quantified after treatment with Proteinase K at different time points. Aptamers or antibodies against CD19 or CD20 were incubated with Raji or Ramos cells, followed by washing, treatment with varying concentrations of Proteinase K, and incubation at 37°C for 10 minutes. A significant decrease in CD19 aptamer/antibody fluorescence intensity was observed after treatment with Proteinase K compared to no treatment. We observed that all cells were partially digested and that no signal was detected after optimized Proteinase K treatment, thus providing an effective platform by which to assess the internalization of dimeric anti-CD19 aptamers (**Fig. S9**). Additionally, the stability of the aptamers was verified by incubation for 24 hours in 10% FBS complete medium at 37°C in a 5% CO₂ incubator. Gel electrophoresis confirmed that the aptamers remained intact and were not degraded by serum proteinases, ensuring their functionality under physiological conditions (**Fig. S10**).

Having demonstrated an effective internalization platform, the internalization of three aptamers, WB15/15.CD19.1_3S, WB17/17.CD19.1_3S, and WB15/17.CD19.1_3S, was assessed using an anti-CD19 mAb as a positive control and a mutated dimeric aptamer as a negative control. The aptamers and controls were incubated on ice for 50 minutes, and their binding was analyzed using flow cytometry. The samples were then incubated at 37°C for 0.5, 1, 3, 6, 12, 24, and 48 hours, followed by Proteinase K treatment. The internalized signal was monitored at each time point. Interestingly, neither the anti-CD19 antibody nor the three tested dimeric aptamers internalized into Burkitt’s lymphoma Raji cells (percent internalization ∼<1%) when compared to that in Ramos cells (**Fig. 2; Fig. S11**). The anti-CD19 antibody rapidly internalized into Ramos cells within 30 minutes, aligning with previously observed rapid internalization of the anti-CD19 mAb (**Fig. 2A; Fig. S11A**). All aptamer constructs, however, showed slow internalization until 6 hours. Importantly, the choice of fluorescent label (e.g., APC, CY3, or FAM) did not affect internalization, as demonstrated by the lack of signal in the isotype and random DNA controls (**Fig. S11**). The observed internalization was driven specifically by the binding of the anti-CD19 antibody and the dimeric CD19 aptamers, and not by the fluorescent dye (**Fig. S11**). Therefore, despite binding to the same target, the internalization kinetics of the dimeric aptamers against CD19 are much slower than that of the anti-CD19 antibodies (**Fig. 2C; E; G; Fig. S11C; S11E; S11G**). Finally, we observed potential receptor recycling. Beyond 36 hours, the percentage of internalized aptamers dropped by ∼50%, and a similar pattern was observed for the anti-CD19 mAb (**Fig. 2; Fig. S11**).

**Figure 2:**
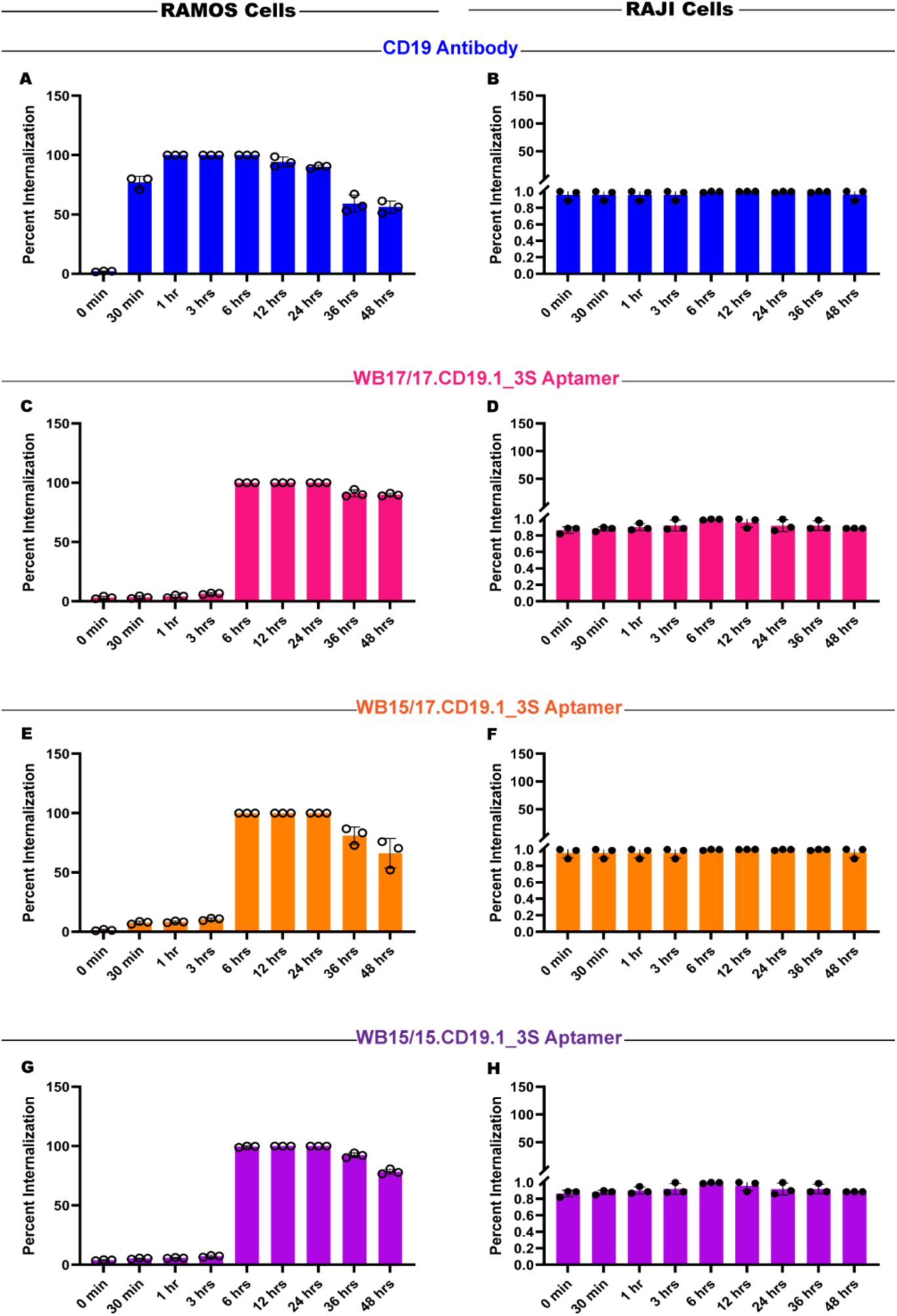
Internalization Kinetics of Dimeric CD19 Aptamers in B-cell Lines. Internalization studies of CD19 Antibody and Bivalent CD19 Aptamers in Ramos and Raji Cells. (A–H) Time-course analysis of CD19 antibody and bivalent CD19 aptamer internalization in Ramos cells (A, C, E, and G: CD21-negative) and Raji cells (B, D, F, and H: CD21-positive). (A-B) demonstrate internalization of APC-CD19 antibody in Ramos (A) and Raji (B) cells. In Ramos cells (A), CD19 antibody shows efficient internalization over 48 hours, while in Raji cells (B), CD21 expression inhibits internalization. (C-D) show internalization of bivalent CD19 aptamer WB17/17.CD19.1_3S in Ramos (C) and Raji (D) cells. WB17/17.CD19.1_3S shows robust internalization in Ramos cells (C), but reduced uptake in Raji cells owing to CD21-mediated blocking. (E-F) illustrate internalization of bivalent CD19 aptamer WB15/17.CD19.1_3S in Ramos (E) and Raji (F) cells. The aptamer demonstrates high internalization efficiency in Ramos cells (E), but not CD21-positive Raji cells (F). Panels G and H: Internalization of bivalent CD19 aptamer WB15/15.CD19.1_3S in Ramos (G) and Raji (H) cells. Similar to other bivalent aptamers, WB15/15.CD19.1_3S internalizes efficiently in CD21-negative Ramos cells (G), but CD21 expression in Raji cells (H) significantly blocks its internalization. Data are expressed as the percentage of internalization calculated as 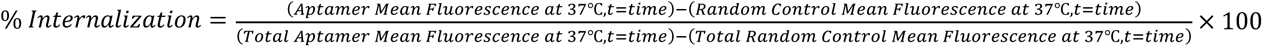. Each bar represents mean ± standard deviation from three independent experiments.

We asked why neither dimeric aptamers nor anti-CD19 antibody internalized into Raji cells. We noted previous studies showing that co-expression of CD19 with the receptor CD21 inhibits the internalization of the CD19 antigen.^26–28^ Following this line of evidence, we assessed whether CD21 is expressed on Raji, Toledo, and Ramos cells. Consistent with findings by others, we observed that CD19 and CD21 are, indeed, highly expressed on the two cell lines that showed no internalization of either CD19 aptamers or antibody (**Fig. S12A-D for Raji, S12F-G for Toledo, S12I-L for Ramos**). Based on this information, we used flow cytometry and confocal microscopy to determine if dimeric aptamers against CD19 colocalize with CD21 on Raji cells. As expected, anti-CD19 mAb, anti-CD21 antibody and the dimeric CD19 aptamers colocalized with CD21, confirming that the presence of CD21 prevents the internalization of the CD19 aptamers (**Fig. 3A-B, 3C-E, 3F-G, 3H-J**).

**Figure 3:**
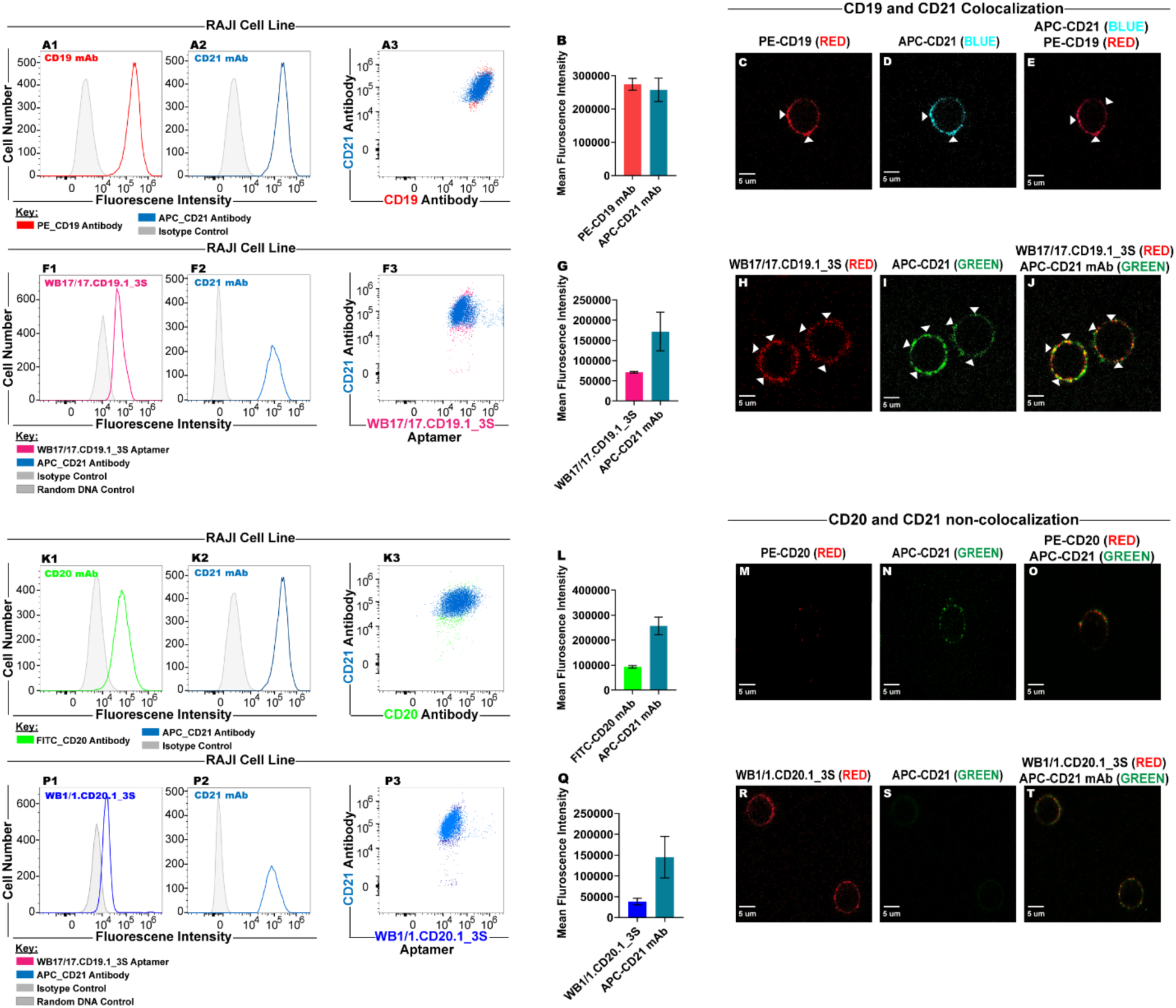
Colocalization of CD19 Dimeric Aptamers and CD21 in Raji cells. Colocalization of CD19 and CD21 was confirmed by flow cytometry and confocal microscopy, whereas CD20 does not co-localize with CD21. (A) demonstrates flow cytometry histograms showing CD19 (A1: PE-CD19 mAb, Red) and CD21 (A2: APC-CD21 mAb, Light Blue) fluorescence intensity in Raji cells. (A3) Bi-parametric dot plot confirms colocalization of CD19 and CD21 on the same population of cells. (B) shows bar graph quantifying mean fluorescence intensity of CD19 and CD21, highlighting their robust expression on Raji cells. (C–E) Confocal microscopy images showing CD19 (C: PE-CD19, Red) colocalized with CD21 (D: APC-CD21, Light Blue) on the surface of Raji cells. Arrowheads indicate regions of colocalization (D). Panels F1–F3 illustrate flow cytometry analysis of bivalent CD19 aptamer WB17/17.CD19.1_3S (F1: Pink) binding to Raji cells and its colocalization with CD21 (F2: APC-CD21 mAb, Light Blue). (F3) Dot plot shows overlapping signals, confirming aptamer-CD21 interaction. (G) demonstrates the bar graph of mean fluorescence intensity of bivalent CD19 aptamer (WB17/17.CD19.1_3S) and CD21 antibody. (H–J) present confocal microscopy images showing WB17/17.CD19.1_3S (H: Red) colocalized with CD21 (I: Green) on Raji cells. Arrowheads highlight colocalized regions (J). (K1–K3) Flow cytometry analysis showing CD20 (K1: FITC-CD20 mAb, Green) and CD21 (K2: APC-CD21 mAb, Light Blue) fluorescence intensity in Raji cells. (K3) Dot plot shows no significant overlap between CD20 and CD21, indicating no colocalization. (L) Bar graph showing fluorescence intensity of CD20 and CD21. (M–O) Confocal microscopy images showing no colocalization between CD20 (M: Red) and CD21 (N: Green) on Raji cells (O). (P1–P3) Flow cytometry analysis of bivalent CD20 aptamer WB1/1.CD20.1_3S (P1: Blue) and CD21 (P2: APC-CD21 mAb, Light Blue). (P3) Dot plot confirms the absence of colocalization on the same population of cells. Panel Q: Bar graph of fluorescence intensity for CD20 aptamer and CD21. (R–T) Confocal microscopy images showing that WB1/1.CD20.1_3S (R: Red) and CD21 (S: Green) do not co-localize on Raji cells (T). Scale bars = 5 μm. Mean fluorescence intensity was calculated using the formula: Mean fluorescence intensity=Aptamer Mean Fluorescence − Random DNA Mean Fluorescence for aptamer. As for antibody it was calculated using the formula: Mean fluorescence intensity=Antibody Mean Fluorescence − Isotype Control Mean Fluorescence. The aptamer/random and antibody/isotype mean fluorescence values corresponds to the mean fluorescence observed in their respective histograms. Data represents mean ± standard deviation from three independent experiments. Each bar represents mean ± standard deviation from three independent experiments.

We extended our study to diffuse large B cell lymphoma (DLBCL) cells, which known grow rapidly and uncontrollably, specifically the germinal center B cell-type (DLBCL-GCB) cell lines OCI-LY7 and HBL-1. Both OCI-LY7 and HBL-1 express high levels of CD19 and CD20, but not CD21 (**Fig. 4C-D, G-H**). The three tested dimeric aptamers against CD19 showed specific binding to both cell lines (**Fig. 4A1-A3, B, E1-E3, F**). The dimeric aptamer WB1/1.CD20.1_3S also showed specific binding to both cell lines (**Fig. 4A4, B, E4, F**). Optimization by Proteinase K assay revealed that these cells require a higher dose of Proteinase K for effective digestion (**Fig. S13**). Internalization was assessed at the 24-hour time point, and we observed an internalization pattern similar to that seen in Ramos cells, suggesting again that the absence of CD21 facilitates the internalization of dimeric aptamers (**Fig. 4I-L**). Confocal microscopy further confirmed that both dimeric aptamers and antibodies internalized into Ramos and OCI-LY7 cells (**Fig. 5A-I for Ramos, 5K-S for OCl-LY7**).

**Figure 4:**
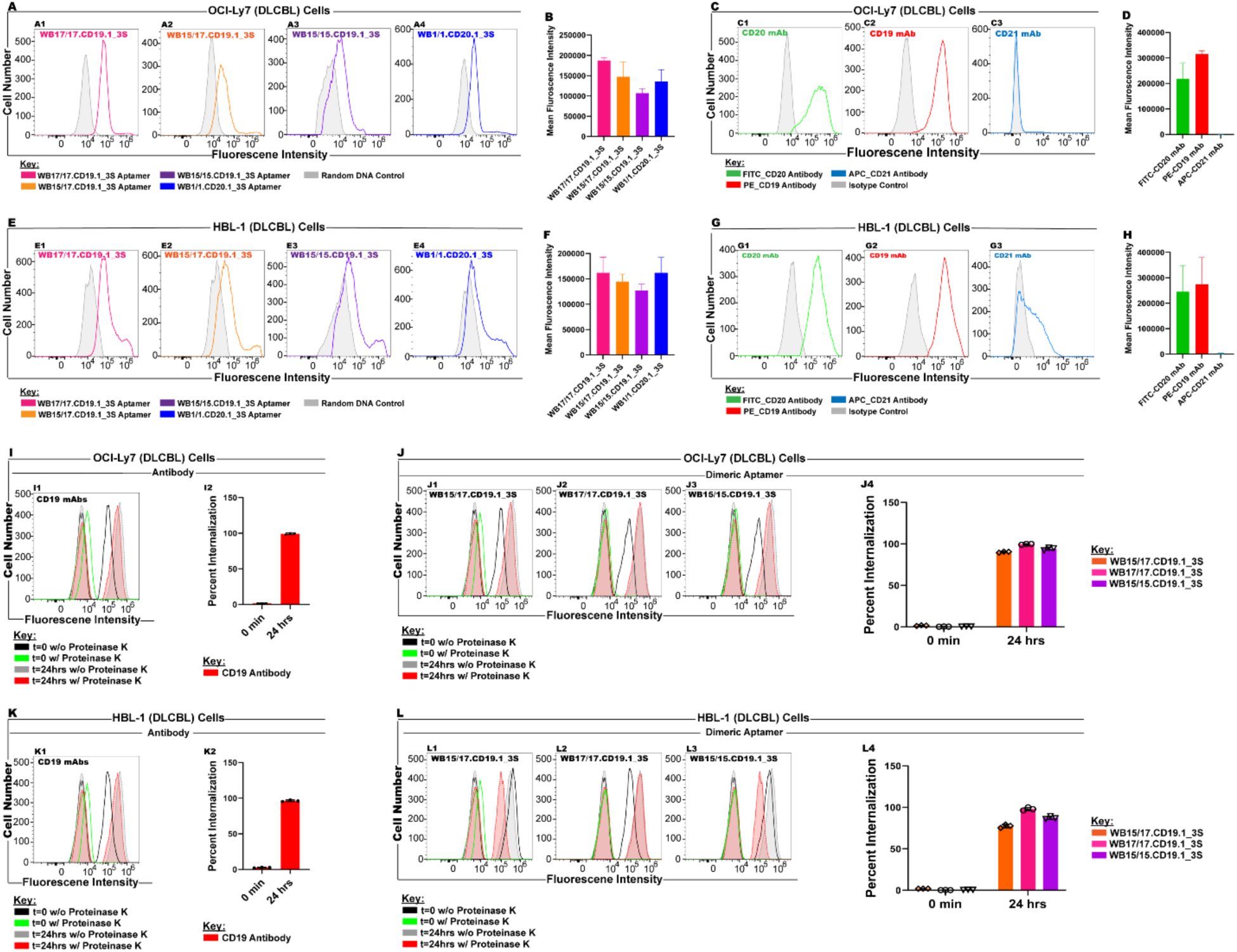
Analysis of CD19, CD20, and CD21 Expression and Internalization in DLBCL Cell Lines. The expression of CD19, CD20, and CD21, the binding activity of bivalent CD19 aptamers, and the internalization dynamics of CD19 antibody and bivalent aptamers in OCl-LY7 and HBL-1 (DLBCL) cells. Antibody staining for CD19, CD20, and CD21 expression. (C; G) demonstrate flow cytometry histograms showing the expression of CD19 (C2: Red), CD20 (C1: Green), and CD21 (C3: Light Blue) on OCI-Ly7 (C), whereas HBL-1 (G) cells express CD19 (G2: Red), CD20 (G1: Green), and CD21 (G3: Light Blue). Isotype controls (gray) confirm specific binding. (D; H) Bar graphs showing mean fluorescence intensities, confirming robust expression of CD19 and CD20, but the absence of CD21 expression in both cell lines. Binding assay with bivalent CD19 aptamers (A1–A4, E1–E4). Fluorescence intensity histograms of bivalent CD19 aptamers (WB17/17.CD19.1_3S, WB15/17.CD19.1_3S, and WB15/15.CD19.1_3S) compared to random DNA control in OCI-Ly7 (A1–A4) and HBL-1 (E1–E4) cells. (B; F) illustrate bar graphs of mean fluorescence intensities showing high binding specificity of bivalent aptamers compared to controls. Internalization of CD19 antibody with and without Proteinase K (I, K). Flow cytometry histograms (L1) and bar graphs (L2) showing internalization dynamics of CD19 antibody in OCI-Ly7 (I), whereas flow cytometry histograms (K1) and bar graphs (K2) show internalization dynamics of CD19 antibody in HBL-1 (K) cells. Internalization was analyzed at 0 hour and 24 hours in the presence and absence of Proteinase K. Data reveal a significant reduction in surface fluorescence intensity after Proteinase K treatment, confirming internalization. Internalization of bivalent CD19 aptamers under the same conditions (J, L). Fluorescence intensity histograms (J1-J3, L1-L3) and bar graphs (J4 and L4) show the internalization of bivalent CD19 aptamers (WB17/17.CD19.1_3S (J2, L2), WB15/17.CD19.1_3S (J1, L1), and WB15/15.CD19.1_3S (J3, L3)) in OCI-Ly7 (J) and HBL-1 (L) cells. Internalization is measured at 0 hour and 24 hours with and without Proteinase K, demonstrating the effective uptake of bivalent aptamers. Data are expressed as the percentage of internalization calculated as 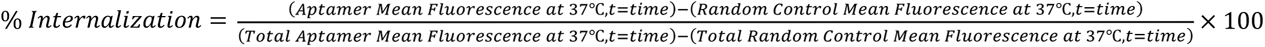 fluorescence intensity was calculated using the formula. Mean fluorescence intensity=Aptamer Mean Fluorescence − Random DNA Mean Fluorescence for aptamer. As for antibody it was calculated using the formula: Mean fluorescence intensity=Antibody Mean Fluorescence − Isotype Control Mean Fluorescence. The aptamer/random and antibody/isotype mean fluorescence values corresponds to the mean fluorescence observed in their respective histograms. Each bar represents mean ± standard deviation from three independent experiments. Data are presented as mean ± standard deviation from three independent experiments.

**Figure 5:**
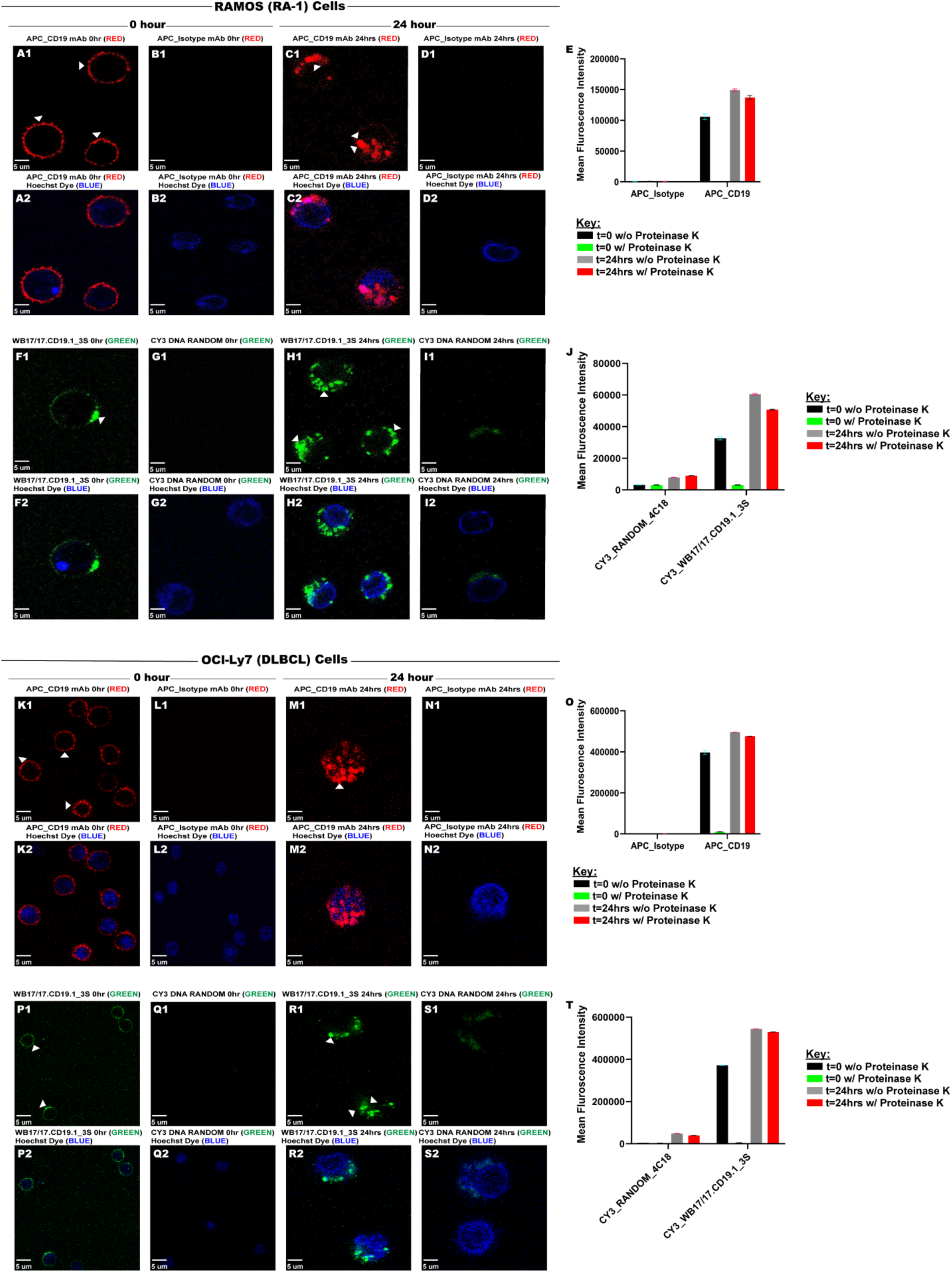
Visualization of CD19 antibody and Bivalent CD19 Aptamer Internalization by Confocal Microscopy. Internalization Assay of CD19 Antibody and Bivalent CD19 Aptamer in Ramos and OCI-Ly7 Cells Visualized by Confocal Microscopy. Internalization of CD19 Antibody in Ramos Cells (A-D). Confocal images showing surface-bound APC-CD19 antibody (A1-A2: Red) at 0 hour with and without Hoechst nuclear staining (A2: Blue). (B1–B2) demonstrate the isotype control at 0 hour. Panels C1-C2 show the internalization of APC-CD19 after 24 hours (C1-C2: Red) with and without Hoechst nuclear staining (C2: Blue). (E) Bar graph quantifying the mean fluorescence intensity of APC-CD19 antibody with and without Proteinase K treatment at 0 and 24 hours. Internalization of Bivalent CD19 Aptamer WB17/17.CD19.1_3S in Ramos Cells (F-I). Initial binding of WB17/17.CD19.1_3S (F1-F2: Green) at 0 hours, as shown with Hoechst-stained nuclei (F2: Blue). (H1–H2) demonstrate internalization of WB17/17.CD19.1_3S at 24 hours with and without Hoechst nuclear staining (H2: Blue). (J) shows the bar graph quantifying the mean fluorescence intensity of WB17/17.CD19.1_3S with and without Proteinase K treatment at 0 and 24 hours. Internalization of CD19 Antibody in OCI-Ly7 Cells (K-N) Surface-bound APC-CD19 antibody at 0 hours (K1-K2). (M1–M2) illustrate reduced surface-bound fluorescence at 24 hours, consistent with internalization. After 24 hours, CD19 antibody is completely uptaken on OCl-Ly7 cells. (O) Bar graph quantifying the mean fluorescence intensity of APC-CD19 antibody with and without Proteinase K treatment at 0 and 24 hours. Internalization of Bivalent CD19 Aptamer WB17/17.CD19.1_3S in OCI-Ly7 Cells (P-S). Confocal images showing WB17/17.CD19.1_3S (P1: Green) binding at 0 hours with and without Hoechst nuclear staining (P2: Blue). (R1–R2) highlight internalized WB17/17.CD19.1_3S at 24 hours with Hoechst nuclear staining (R2). Panel T shows the bar graph quantifying the mean fluorescence intensity of WB17/17.CD19.1_3S with and without Proteinase K treatment at 0 and 24 hours. Scale Bars: 5 μm. Data represent mean ± standard deviation from three independent experiments. Each bar represents mean ± standard deviation from three independent experiments.

### Dimeric anti-CD20 aptamer does not induce intracellular calcium release in DLBCL cell lines

We investigated the potential of aptamers to induce intracellular calcium release by using three diffuse large B-cell lymphoma (DLBCL) cell lines: OCI-LY7, HBL-1, and Karpas 422. Using random DNA control, WB15/15.CD19.1_3S, WB17/17.CD19.1_3S, and WB1/1.CD20.1_3S aptamers, we monitored intracellular calcium levels using the fluorescent calcium indicator Fluo-4 AM. The fluorescence intensity was recorded at 535 nm following excitation at 485 nm. Across all three DLBCL cell lines, no significant changes (ns) were observed in calcium mobilization in response to the aptamers when compared to the random DNA control (**Fig. S14A-C**). This finding suggests that the aptamers do not modulate calcium signaling under the conditions tested.

## Discussion

Expressed in late pre-B lymphocytes, CD20 is a non-glycosylated protein that belongs to the membrane-spanning 4-domains (MS4) protein family.^29, 30^ It plays a crucial role in B-cell activation, differentiation, and calcium ion regulation.^31^ Owing to its restricted expression on B cells and absence on stem cells or plasma cells, CD20 has become a key therapeutic target in the treatment of B-cell malignancies^32^. Although the actual biological role of CD20 remains unclear, Rituximab, an antibody developed against CD20, is one of the major therapeutic modalities in the treatment of B-cell malignancies and several autoimmune diseases, and over a million doses of Rituximab have been administered to treat B-cell-related diseases.^3^ Thus, CD20 remains a crucial target for the development of novel therapeutic strategies, including bispecific antibodies and antibody-drug conjugates.^33^ The antigen CD19, also known as B4, on the other hand, is a type 1 transmembrane glycoprotein that belongs to immunoglobulin superfamily (IgSF) that is specifically expressed in normal and neoplastic B cells and follicular dendritic cells (FDCs).^34^ CD19 is expressed in pre-B-cells and mature B-cells, and it is categorized as one of the essential biomarkers in B-cells.^35^ It plays a key role in B-cell receptor (BCR) signaling, as well as B-cell activation, differentiation, and survival.^36, 37^ CD19 is also widely expressed in most B-cell malignancies, including acute lymphoblastic leukemia (ALL) and non-Hodgkin’s lymphoma, making it an important therapeutic target.^38–40^ Herein, we developed two synthetic, highly specific dimeric aptamers against CD19 and CD20 expressed on B-cells.

We previously introduced dimeric aptamers against Membrane-Bound Immunoglobulin M (mIgM) and TCR-CD3 and demonstrated that systematic dimerization can lead to aptamer assemblies with higher specificity and affinity.^17, 18^ In this study, we employed a similar strategy that first involved truncation and then dimerization of CD19 and CD20 aptamers, followed by an evaluation of their specificity, avidity, and biochemical functionality.^41^ The rationale for dimerization is multifaceted. First, dimerization results in a higher local concentration of the aptamers, effectively improving their affinities. Second, it increases the molecular weight of the aptamers, potentially enhancing their circulation time. Third, tethering aptamers with PEG linkers can improve nuclease resistance. To achieve dimerization, we utilized PEG as the linker to introduce linear assemblies of dimeric aptamers. Linear assemblies offer significant advantages over spherical or single-layer two-dimensional assemblies because they can more readily adopt functional conformations with minimal steric hindrance. The flexibility of PEG linkers further minimizes steric hindrance and allows for maximum interaction with the target.

Our findings confirm that careful optimization of linker length is critical for developing functional linear dimeric aptamers. Interestingly, when linker lengths are shorter than ∼3.96 nm, both assemblies exhibit behavior similar to that of monomeric aptamers. similarly, when linker lengths are excessively long, they still exhibit monomer-like behavior. For both CD19 and CD20 dimeric aptamers, the optimal linker length was determined to be 3.96 nm. This suggests that larger aptamers with 65 nucleotides may require shorter linkers compared to smaller aptamers containing 35 to 45 nucleotides. The strategic dimerization of CD19 and CD20 aptamers using optimized PEG linkers boosts their biochemical functionality. This approach highlights the importance of linker design in developing aptamer-based assemblies with improved specificity and affinity.

We previously demonstrated that aptamers identified using LIGS exhibit biochemical characteristics similar to the secondary ligand used for selective elution. For instance, the dimeric aptamer R1.2 displayed specificity toward both soluble and membrane-bound IgM akin to that of anti-IgM antibodies.^18^ Similarly, the dimeric anti-CD3 aptamer activated Jurkat.E6 cells.^17^ In this study, we found that, similar to anti-CD19 antibody, the dimeric anti-CD19 aptamer selectively internalizes into B-cells based on CD21 expression levels. Thus, the dimeric aptamers introduced here exhibit a selective internalization pattern similar to that of the B4 antibody. Specifically, like B4, the internalization of dimeric anti-CD19 aptamers is negatively correlated with CD21 expression levels. For instance, our findings showed that all tested dimeric anti-CD19 aptamers internalized into Ramos, OCl-LY7, and HBL-1 cells, which do not express high levels of CD21. In contrast, neither aptamers nor B4 antibody internalized into Raji cells, which are known to express high levels of CD21. Notably, approximately 30% of DLBCL cases lack CD21 expression, while another 30% exhibit low CD21 levels. These observations suggest that the selective uptake of anti-CD19 aptamers could be potentially useful for engineering targeted therapeutics against DLBCL. Furthermore, the lack of uptake by dimeric anti-CD20 aptamers and the slow uptake of anti-CD19 aptamers present opportunities to develop synthetic bispecific agents for therapeutic applications.

Several such aptamers are currently in clinical trials. Antisense oligonucleotides (ASOs) and small interfering RNAs (siRNAs) are both used to target specific RNA sequences as drug targets, and both can target diffuse large B-cell lymphoma. Similarly, aptamers could be generated that selectively internalize into cells, thus providing a powerful avenue to develop nucleic acid drugs against DLBCL, especially if both drug and targeting agent are made from nucleic acids. For example, the delivery of antisense therapeutics, or mRNA, could be facilitated more easily since both targeting agent and drug are made from nucleic acids, simplifying large-scale synthesis and purification of the aptamer-drug conjugate. In conclusion, the nucleic acid dimeric aptamers herein reported could be developed into targeted therapies against DLBCL and B-cell-related diseases.

## Materials and Methods

### Cell Lines

Toledo (non-Hodgkin’s B cell lymphoma; CRL-2631), Ramos (Clone RA 1, Burkitt’s lymphoma; CRL-1596), Daudi (Burkitt’s lymphoma; CCL-213) and Jurkat (Clone E6-1, acute T cell leukemia; TIB-152) were purchased from American Type Culture Collection (ATCC, Manassas, VA, USA). BJAB (Burkitt’s lymphoma), SKLY-16 (B cell lymphoma), and Raji (Burkitt’s lymphoma) were generous gifts from the David Scheinberg Lab and Huse Lab, Memorial Sloan Kettering Cancer Center. CA46 CD19- and CD20-knockout cells, generated via CRISPR-Cas9, were purchased from Synthego (Redwood City, CA, USA). Toledo cell cultures were maintained in RPMI-1640 medium (+25 mM HEPES, +L-Glutamine, Catalog no. SH30255.01, HyClone) supplemented with 20% fetal bovine serum (Dialyzed 12–14kD MW, Heat Inactivated at 56°C, Catalog no. S12850H, R&D Systems), 100 units/mL penicillin-streptomycin (10,000 I.U./mL Penicillin, 10,000μg/mL Streptomycin, Catalog no. 30-001-Cl, Corning) and 1% MEM non-essential amino acids (100X, Catalog no. 11140-050, Gibco). Ramos, Daudi, BJAB, SKLY-16, Raji, CA46KO, and Jurkat cell cultures were maintained in RPMI-1640 medium (+25 mM HEPES, +L-Glutamine, HyClone) supplemented with 10% fetal bovine serum (Dialyzed 12–14kD, Heat Inactivated), 100 units/mL penicillin-streptomycin (10,000 IU Penicillin, 10,000μg/mL Streptomycin, Corning), and 1% MEM Non-Essential Amino Acids (100X, Gibco). All cell lines were routinely evaluated on a flow cytometer (Cytek, Northern Light-2000) for the expression of their cluster of differentiation (CD) biomarkers using respective monoclonal antibodies to authenticate the cell line. OCI-Ly7 and HBL1 diffuse large B-cell lymphoma (DLBCL) cell lines were kindly provided by Dr. Leandro Cerchietti’s Laboratory at Weill Cornell Medicine.

### Antibodies

PE-conjugated CD19 anti-human monoclonal antibody (mouse, isotype IgG1, Clone 4G7, Catalog no. PE-65|97, Proteintech), PE-conjugated CD19 mouse anti-human (isotype IgG1, κ, Clone SJ25C1, Catalog no. 12-0198-42, Invitrogen), PE-conjugated CD20 anti-human (mouse, isotype IgG2b, κ, Clone 2H7, Catalog no. 302305, BioLegend), PE Mouse IgG1, κ, isotype control (CloneMOPC-21, Catalog no. 556650, BD Pharmingen), FITC-conjugated CD20 anti-human monoclonal antibody (mouse, isotype IgG2b, Clone 2H7, Catalog no. 35-0209-T100), FITC-conjugated CD19 mouse anti-human (isotype IgG1, κ, Clone SJ25C1, Catalog no. 363008, BioLegend), Monoclonal Rabbit IgG Alexa Fluor 488 (Clone 60024B; Catalog no. IC1051G, R&D Systems, MN), APC Mouse IgG1, κ, Isotype Ctrl (FC) (Clone MOPC-21; Catalog no. 400122), APC-conjugated CD19 anti-human (mouse, isotype IgG1, κ, Clone SJ25C1, Catalog no. 17-0198-42, Invitrogen), APC-conjugated mouse anti-human CD20 (isotype IgG2b, κ, Clone 2H7, Catalog no. 559776, BD Pharmingen), and APC-conjugated CD21 mouse anti-human (isotype IgG2a κ, Clone HB5, Catalog no. 17-0219-42, Invitrogen) were used for routine flow cytometry analysis.

### Cell Suspension Buffer (CSB)

All *in vitro* experiments were performed using a Cell Suspension Buffer (CSB) formulated with HyClone RPMI-1640 medium (+25mM HEPES, +L-Glutamine) containing 200 mg/L tRNA from Baker’s yeast (Catalog no. J61215.ME, ThermoScientific), 2g/L Bovine Serum Albumin (BSA, CAS 232-936-2, Fisher Scientific), and 10 mg/L Salmon Sperm DNA solution (Catalog no. 15632011, Invitrogen). The wash buffer used was HyClone RPMI-1640 medium (+25mM HEPES, +L-Glutamine). The tRNA, BSA, and salmon sperm DNA were added to block nonspecific binding sites on the cell surface.

### Antibody Staining Assay

Cells were prepared by washing three times with 3 mL of RPMI-1640 medium, followed by centrifugation at 1000 rpm for 3 minutes. The cell pellet was resuspended in Cell Staining Buffer (CSB) to a final volume of 100 µL, containing 2.0 × 10⁵ cells, and incubated at 25°C for 15 minutes. After incubation, the 100 µL cell suspension was transferred into flow cytometry tubes and placed on ice. Antibodies were added at a final concentration of 0.03 µg per tube, and the samples were incubated on ice for 20–30 minutes to allow for staining. Subsequently, the cells were washed with 3 mL of RPMI-1640 medium, resuspended in 250 µL of RPMI-1640, and analyzed using a Cytek NL-2000 flow cytometer. Data acquisition and analysis were performed using FlowJo software (version 10.10.0).

### DNA Synthesis

All DNA reagents needed for DNA synthesis were purchased from Glen Research. The monomeric WB1.CD20, WB1.CD20.1, WB1.CD20.2, WB1.CD20.3, WB15.CD19, WB15.CD19.1, WB15.CD19.2, WB17.CD19, WB17.CD19.1, and WB17.CD19.2 molecules were chemically synthesized by attaching a fluorophore at the 5’-end using standard solid-phase phosphoramidite chemistry on an ABI394 DNA/RNA Synthesizer (Biolytic) using a 1 µmol scale. The dimeric molecules were chemically synthesized by attaching a fluorophore at the 5’-end using standard solid-phase phosphoramidite chemistry on a ABI394 DNA/RNA Synthesizer using a 1 µmol scale, and a Spacer Phosphoramidite 9 (Catalog no. 10-1909-90, Glen Research) was used to tether the two monomeric molecules. One, two, three, and four spacers were used for WB1/1.CD20.1_1S, WB15/15.CD19.1_1S, WB15/17.CD19.1_1S, WB17/17.CD19.1_1S, WB1/1.CD20.1_2S, WB15/15.CD19.1_2S, WB15/17.CD19.1_2S, WB17/17.CD19.1_2S, WB1/1.CD20.1_3S, WB15/15.CD19.1_3S, WB15/17.CD19.1_3S, WB17/17.CD19.1_3S, WB1/1.CD20.1_4S, and WB17/17.CD19.1_4S, respectively. The completed DNA sequences were deprotected according to the base modification employed and purified using High-Performance Liquid Chromatography (HPLC, Agilent) equipped with a C-18 reversed phase column (Phenomenex/Waters). Monomeric and dimeric DNA Random Controls were purchased from IDT DNA Technologies.

Preparation of Solutions and Folding Conditions

After the purification of monomers (WB1.CD20, WB1.CD20.1, WB1.CD20.2, WB1.CD20.3, WB15.CD19, WB15.CD19.1, WB15.CD19.2, WB17.CD19, WB17.CD19.1, WB17.CD19.2) and dimers (WB1/1.CD20.1_1S, WB15/15.CD19.1_1S, WB15/17.CD19.1_1S, WB17/17.CD19.1_1S, WB1/1.CD20.1_2S, WB15/15.CD19.1_2S, WB15/17.CD19.1_2S, WB17/17.CD19.1_2S, WB1/1.CD20.1_3S, WB15/15.CD19.1_3S, WB15/17.CD19.1_3S, WB17/17.CD19.1_3S, WB1/1.CD20.1_4S, and WB17/17.CD19.1_4S), concentration of the stock solutions was determined by using a UV-Vis Spectrophotometer (Cary 60 UV-Vis, Agilent). Then, 10µM solutions of the monomeric and dimeric molecules were prepared by dilution of the respective stock solutions with RPMI-1640 medium (+25mM HEPES, +L-Glutamine). These 10µM solutions were then used to prepare 2µM, 500nM, and 250nM working solutions by using RPMI-1640 medium (+25mM HEPES, +L-Glutamine). Random controls were prepared in a manner similar to that of each monomeric and dimeric molecule. The folding of random control and anti-CD19 monomeric and dimeric aptamer solutions was done by denaturing at 95°C for 10 minutes and maintaining under dark cover with aluminum at 25°C for 45 minutes. Anti-CD20 monomeric and dimeric aptamers and random control solutions were carried out by heating at 95°C for 10 minutes and maintaining in the dark on ice for 45 minutes. The maximum time of 45 minutes for folding was strictly followed because aptamer binding diminishes if the folded aptamer is kept on ice any longer.

### Determination of Apparent binding affinity at 4°C

Aptamer dilutions and folding were performed as described in the previous section. Apparent binding affinity of monomers and dimers towards target cells were determined by using Raji (1.5 × 10^5^) cells. Both CD20 and CD19 fluorescently-labeled aptamer concentrations were used (500nM to 0.5nM), and cells were incubated in 50 μL of CSB and 50 μL of aptamer solutions in RPMI-1640 for 1 hour on ice. After 1 hour, 3mL of RPMI-1640 medium were added for one wash, and the cells were then reconstituted in 250 μL of RPMI-1640 medium. Aptamer binding towards targets cells was analyzed with flow cytometry for each concentration by recording 5000 events. K_d_ values were obtained by plotting the aptamer’s apparent fraction of bound aptamers, which was calculated as Specific Mean Fluorescence = Aptamer Mean Fluorescence − Random DNA Mean Fluorescence, against the concentration of aptamers with GraphPad Prism software using a nonlinear regression (curve fit) and one site-specific binding.

### Determination of Apparent binding affinityat 37°C

The apparent affinity of target-specific aptamers against Raji cells was determined using a range of aptamer concentrations from 250 nM to 0.25 nM. Both CD19 and CD20 aptamers were prepared as described in the previous section. Cells were prepared before the aptamer binding assays by washing three times with 3 mL RMPI-1640 medium. Each concentration of aptamers was incubated with 1.5 × 10^5^ Raji cells at 37°C for 1 hour in a final volume of 100 μL. Cells were then washed once using 3 mL RPMI-1640 medium, and binding events were analyzed by flow cytometry. Kd values were obtained by plotting the aptamer’s specific mean fluorescence against nanomolar (nM) concentration of aptamers with GraphPad Prism software, using a nonlinear regression (curve fit), one site-specific binding.

### Specificity Assays

Specificity assays were conducted with individual aptamers against HPLC-purified random DNA (monomeric and dimeric) control purchased from IDT DNA Technologies. The specificity of aptamer sequences was evaluated by incubating parents (full length), truncated analogues, and dimeric analogues separately with six different cell lines accompanying the B-cell lines Toledo, Raji, BJAB, Ramos, SKLY-16, OCl-Ly7, HBL-1, and Karpas 422, while the negative cell lines were Jurkat.E6 and CA46 knockdown. These assays were performed by incubating 50 μL of either monomeric or dimeric aptamer (500nM or 250nM) or random control with 2.0 × 10^5^ cells in 50 μL of cell suspension buffer at 37°C for 45 minutes, followed by washing once with 3 mL RPMI-1640 medium. Cells were reconstituted in 250 μL RPMI-1640. Finally, binding was analyzed by flow cytometry by counting 10,000 events. Using FlowJo software (version 10.10.0), we take the mean fluorescence intensity and plot into GraphPad Prism software, one site-specific binding. Three independent experiments against each cell line were performed.

### Fluorescence Microscopy

Toledo cells were washed three times by centrifugation at 1000 rpm for 3 minutes with 3 mL of RPMI-1640 medium between each wash. The cells were then counted, and 2 × 10⁵ cells were resuspended in 100 μL of Cell Staining Buffer (CSB). For binding studies, 2 × 10⁵ cells in 100 μL were incubated with 2 μM of either Cy3-labeled WB1-CD20.1 or WB17-CD19.1 aptamer, or random DNA control, in 100 μL to achieve a final aptamer concentration of 1 μM. Incubations were carried out on ice for 45 minutes. Following this, 0.125 μg of APC-labeled anti-CD20 antibody was added to the tubes incubated with WB1-CD20.1 aptamer or random DNA control. Similarly, 0.125 μg of APC-labeled anti-CD19 antibody was added to the tubes incubated with WB17-CD19.1 aptamer or random DNA control. For controls, 0.125 μg of APC-labeled isotype control antibody was added to tubes incubated with either WB1-CD20.1 or WB17-CD19.1 aptamer. After antibody incubation, binding was disrupted by adding 2 mL of RPMI-1640 medium, followed by centrifugation. The supernatant was decanted, and the cell pellet was resuspended in 500 μL of 1× PBS (Cat. No SH30256.01). The resuspended cells were mixed thoroughly and gently transferred to the center of a glass coverslip in a 35 mm × 10 mm style cell culture dish (Cat. No 430165), ensuring that the cell suspension covered only the glass coverslip by surface tension. Cells were allowed to adhere by gravity sedimentation at room temperature for 30 minutes. Following adhesion, cells were fixed by carefully adding 500 μL of 4% paraformaldehyde (PFA) prepared in 1× PBS to the dish, breaking the surface tension. This resulted in a final PFA concentration of 2%. Fixation was allowed to proceed for 10 minutes at room temperature. After fixation, the fixative was aspirated, and the cells were washed with 1.5 mL of 1× PBS to remove any non-adherent cells. After washing, 1 mL of fresh 1× PBS was added to the sides of the culture dish to maintain hydration. Coverslips with adherent cells were gently lifted from the dish using sharp-tipped tweezers. Excess liquid from the backside of the coverslip was removed with a Kimwipe, and residual liquid on the cell side was gently absorbed by touching the corners of the coverslip with a Kimwipe. Coverslips were mounted cell-side down onto microscope slides pretreated with one drop of Fluoromount-G mounting medium. Slides were allowed to dry for 15 minutes in a light-protected area before imaging with a Nikon TiE inverted microscope (Nikon Inc., Melville, NY). Images were acquired using a Neo sCMOS camera (6.45 μm pixels, 560 MHz, Andor Technology) and a 40X dry objective. This protocol ensured high-quality imaging of Cy3-labeled aptamer and APC-labeled antibody colocalization in Toledo cells.

### Proteinase K Treatment

Following the binding or internalization assay, Raji, Toledo, Ramos, OCl-Ly7, and HBL-1 cells (2.0 × 10⁵ cells per sample) were washed with 2 mL of 1X Phosphate-Buffered Saline (PBS, calcium- and magnesium-free; Catalog no. SH30256.01, HyClone) to remove unbound material. Subsequently, the cells were incubated with 100 µL of proteinase K solution prepared in PBS. For Raji, Toledo, and Ramos cells, 1 mg/mL proteinase K (Catalog no. EO0491, ThermoScientific) was used, while OCl-Ly7 and HBL-1 cells were treated with 3 mg/mL proteinase K. Incubation was performed at 37°C for 10 minutes. After the initial incubation, Raji, Toledo, and Ramos cells were washed with 1 mL of PBS at room temperature and centrifuged at 1000 rpm for 3 minutes; then the supernatant was discarded. The pellet was resuspended in 100 µL of 1 mg/mL proteinase K in PBS and incubated again at 37°C for 10 minutes. Similarly, OCl-Ly7 and HBL-1 cells were washed with 1 mL of PBS at room temperature and centrifuged at 350 rcf for 5 minutes; then the supernatant was discarded. These cells were resuspended in 100 µL of 3 mg/mL proteinase K in PBS and incubated at 37°C for an additional 10 minutes. To terminate proteinase K activity, 100 µL of 20% FBS-containing medium was added to each sample and incubated for 1 minute. The cells were then washed with 3 mL of RPMI-1640 medium. Raji, Toledo, and Ramos cells were centrifuged at 1000 rpm for 3 minutes, while OCl-Ly7 and HBL-1 cells were centrifuged at 350 rcf for 5 minutes. After discarding the supernatant, the cell pellets were resuspended in 250 µL of RPMI-1640 medium. Samples were analyzed using flow cytometry, and data were processed using FlowJo software (version 10.10.0). Graphical representations, including bar graphs, were generated using GraphPad Prism software.

### Internalization Assays

The dimeric aptamer molecules WB15/15.CD19.1_3S, WB15/17.CD19.1_3S, and WB17/17.CD19.1_3S were prepared as previously described and used at a final concentration of 250 nM in a total volume of 100 µL in CSB buffer, alongside dimeric random controls prepared under identical conditions. Raji, Ramos, OCl-Ly7, and HBL-1 cell lines were washed three times with 3 mL RPMI-1640 medium, resuspended in CSB buffer, and incubated at 25°C for 15 minutes before the assay. For the internalization assay, 2.0 × 10⁵ cells per sample were incubated with aptamer solutions in a final volume of 200 µL per well at 4°C or on ice for 1 hour. APC-conjugated CD19 antibody (Clone SJ25C1) and APC-conjugated isotype control were added as positive controls after 20 minutes of incubation at a final concentration of 0.0015 µg in 100 µL. After the initial 1-hour incubation, 100 µL of each sample were transferred into flow cytometry tubes and kept on ice, while the remaining wells received 100 µL of 20% FBS-containing media and were incubated at 37°C in a humidified atmosphere with 5% CO₂ for time points of 1 hour, 3 hours, 6 hours, 12 hours, 24 hours, and 48 hours. For each sample in flow cytometry, tubes were subsequently washed with cold PBS, and proteinase K treatment was performed on half the volume to remove surface-bound aptamers, while the other half was left untreated to assess total aptamer binding. This step ensured discrimination between internalized and surface-bound aptamers. All treatments and washes were conducted as described in prior sections to maintain consistency. This protocol enables precise and reproducible quantification of aptamer internalization dynamics under various experimental conditions.

### Confocal Imaging

Cellular fluorescence imaging was conducted using a Zeiss LSM 880 confocal microscope (Zeiss, USA) equipped with a Plan-Apochromat 63×/1.40 oil immersion DIC objective (63x oil immersion objective). The excitation wavelengths and emission detection ranges were optimized for each fluorophore. APC-conjugated antibodies were imaged using a 633 nm laser line for excitation and a spectral detection range of 660– 690 nm, while Cy3-labeled aptamers and random controls were imaged using a 543 nm laser line for excitation and a spectral detection range of 560–590 nm. To halt internalization, cells were placed on ice immediately following incubation and washed with 2 mL of cold PBS containing BD Hoechst 33342 Solution (Catalog no. 561908, BD Pharmingen) to stain nuclei, followed by a 15-minute incubation on ice. After staining, cells were washed again with 3 mL of PBS without Hoechst 33342 solution. No proteinase K treatment was applied. Before imaging, cells were resuspended in 100 µL of Hanks’ balanced salt solution (HBSS, calcium chloride-free, magnesium chloride-free, and magnesium sulfate-free; Catalog no. 14175-095, Gibco) and transferred onto poly-D-lysine-coated 35 mm glass bottom dishes (Part no. P35GC-1.0-14-C, MatTek Corporation). Cells were allowed to settle on the glass surface for 3 minutes on ice. For colocalization experiments, aptamers and random controls were prepared as described previously, and monoclonal antibodies were added at a final concentration of 0.03 µg. Aptamers (or random controls) and antibodies were simultaneously incubated with cells on ice for 1 hour. Following incubation, cells were washed with 2 mL of cold PBS containing BD Hoechst 33342 solution for nuclear staining and incubated for 15 minutes on ice. A final wash with 3 mL of PBS without Hoechst 33342 solution was performed before imaging. Cells were then resuspended in 100 µL of HBSS and transferred onto poly-D-lysine-coated glass bottom dishes, allowing them to settle for 3 minutes on ice. Images were acquired using Zeiss ZEN 2 software, and post-acquisition analyses, including colocalization assessments, were conducted using ImageJ software.

### Gel electrophoresis

A 3.5% agarose gel was prepared by combining 1.75 g of agarose (Fisher Bioreagents, Agarose Genetic Analysis Grade, Catalog no. BP1356-100) with 50 mL of 1X TBE electrophoresis buffer (Thermo Scientific, Catalog no. B52, 10X TBE). The mixture was heated in a microwave until the agarose was completely dissolved. The molten agarose was poured into a 7 x 7 cm tray and allowed to solidify using the Mini-Sub Cell GT Horizontal Electrophoresis System (Bio-Rad) with casting gates. Electrophoresis was performed at 120 V for 45 minutes. Following electrophoresis, the gel was post-stained with SYBR Gold Nucleic Acid Gel Stain (Invitrogen, Catalog no. S11494) for 45 minutes in the dark. The gel was then rinsed with Milli-Q water to remove excess stain. Samples were prepared at a final concentration of 200 nM in PBS before loading onto the gel. Imaging was performed using a GelDoc Go Gel Imaging System (Bio-Rad) to assess purity and electrophoretic mobility of monovalent and dimeric aptamers.

### Denaturing PAGE Gel Electrophoresis

A 7.5% denaturing PAGE gel was prepared by combining 16.25 mL of Milli-Q water, 2.5 mL of 10× TBE buffer (Thermo Scientific, Catalog no. B52), 6.25 mL of Acrylamide/Bisacrylamide Solution (40%, 37.5:1; Thermo Scientific, Catalog no. J60868.AP), 250 µL of Ammonium Peroxodisulfate (APS; TCI America, Catalog no. A2098), and 12.5 µL of N,N,N’,N’-Tetramethylethylenediamine (TEMED; molecular biology reagent, MP Biomedicals, Catalog no. 194019) to a final volume of 25 mL. The mixture was used to cast two gels in the SureCast™ Gel Handcast Station (Invitrogen by Thermo Fisher Scientific, Catalog no. HC1000) and allowed to polymerize for 15–20 minutes. While the gels polymerized, a degradation buffer was prepared by mixing 50 mL of 10× TBE, 5 mL of 10% Sodium Dodecyl Sulfate (UltraPure™ SDS Solution, Thermo Fisher Scientific, Catalog no. 15553027), and 445 mL of Milli-Q water. Once polymerized, the gels were pre-run at 250 V for 30 minutes in the Invitrogen™ XCell SureLock™ Mini-Cell system (Fisher Scientific, Catalog no. EI0001). Following the pre-run, the wells were washed with buffer to ensure cleanliness before sample loading. Sample Preparation and Gel Loading: Samples were prepared by incubating aptamers with cells in 20% FBS media (1:1 ratio, 10% FBS media) for 24 hours in wells of a 96-well plate. After incubation, the samples were transferred to 1.5 mL centrifuge tubes and denatured at 95°C for 10 minutes. Following denaturation, the samples were centrifuged to spin down the cells into pellets. Only the supernatant containing the aptamers was collected for further analysis, while the cell pellet was discarded. A 1:1 ratio of denaturing loading buffer (without dye) was added to the supernatant. For gel loading, 20 µL of the prepared supernatant sample and 10 µL of DNA ladder were loaded into the wells. The gel ran at 200 V for 20 minutes.

### Calcium Release Assays

Cells were incubated with 1 μM Fluo-4 AM (F14217, Invitrogen) for 30 minutes at 37°C. The fluorescent indicator was then washed out with fresh, warm media, and the cells were allowed to recover for an additional 30 minutes at 37°C. After washing, the cells were seeded into wells of a 96-well plate at a density of 50,000 cells per well in 100 μL. Treatments were injected by adding 50 μL of solutions at three times their final concentration (final aptamer concentration: 250 nM). Fluorescence emissions from the 96 wells were monitored simultaneously at an emission wavelength of 535 nm after excitation at 485 nm (F488). Fluorescence data were collected and analyzed offline.

### Mass Spectrometry

For standard nucleotide analysis by ESI-MS MW confirmation, samples are analyzed with a Thermo Scientific Vanquish UHPLC and Thermo Scientific LTQ-XL Ion Trap mass spectrometer. Samples are loaded onto a small trap column in buffer A (0.075% HFIPA hexafluoro-2-propanol/ 0.0375% DIEA N,N-Diisopropylethylamine in water) and eluted in buffer B (0.075% HFIPA hexafluoro-2-propanol/0.0375% DIEA N,N-Diisopropylethylamine 65:35 Acetonitrile:Water). Mass spectra were recorded by Novatia, LLC.

