## Supporting Information for "The biochemical function of bivalent aptamer assemblies against B-cell markers CD19 and CD20"

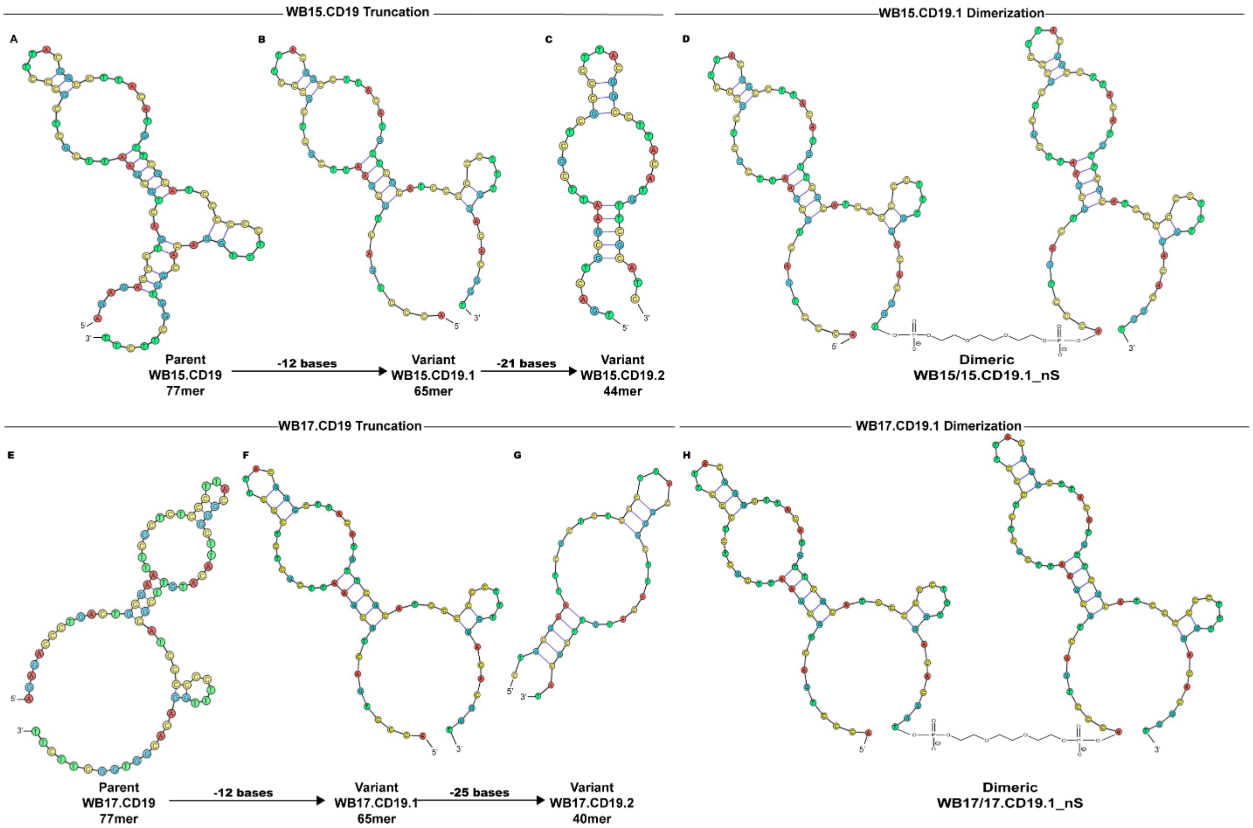


**Figure S1: Predicted Secondary Structures of CD19 Aptamers and their Truncated and Dimerized Variants**. Predicted secondary structures of WB15.CD19 and WB17.CD19 aptamers showing truncation and dimerization for engineering enhanced functionality. (A–H) Aptamer structures were predicted using m-fold software and NUPACK, visualized with Ribosketch. The figure highlights the stepwise truncation from parental aptamers to optimized truncation variants and further to engineered dimeric forms. WB15.CD19 Truncation and Dimerization: (A) The parental WB15.CD19 aptamer (77mer) exhibits a complex multi-loop structure critical for efficient CD19 binding. (B) The first truncation variant, WB15.CD19.1 (65mer), retains key structural motifs. Although it eliminates 12 bases for structural simplification, such elimination dramatically reduces affinity. (C) WB15.CD19.2 (44mer) truncates an additional 21 bases, resulting in no affinity. (D) The dimeric WB15/15.CD19.1_nS, which combines two WB15.CD19.1 units connected by a 9-PEG linker, is designed to boost target binding through multivalent interactions. WB17.CD19 Truncation and Dimerization: (E) The parental WB17.CD19 aptamer (77mer) exhibits a stable structure with prominent loops that ensure high affinity to CD19. (F) The first truncation variant, WB17.CD19.1 (65mer), eliminates 12 bases, while preserving critical binding domains. (G) WB17.CD19.2 (40mer) removes an additional 25 bases, producing a minimal functional unit. However, it loses its affinity. (H) Dimeric WB17/17.CD19.1_nS incorporates two WB17.CD19.1 units linked by a 9-PEG spacer, enabling simultaneous binding to multiple CD19 molecules for increased affinity and stability. Key Insights: Truncation simplifies aptamer structures while maintaining or modulating binding affinity. Dimerization leverages are the best monovalent truncation variants to achieve multivalent binding, offering improved functionality and potential applications in targeted therapeutics.

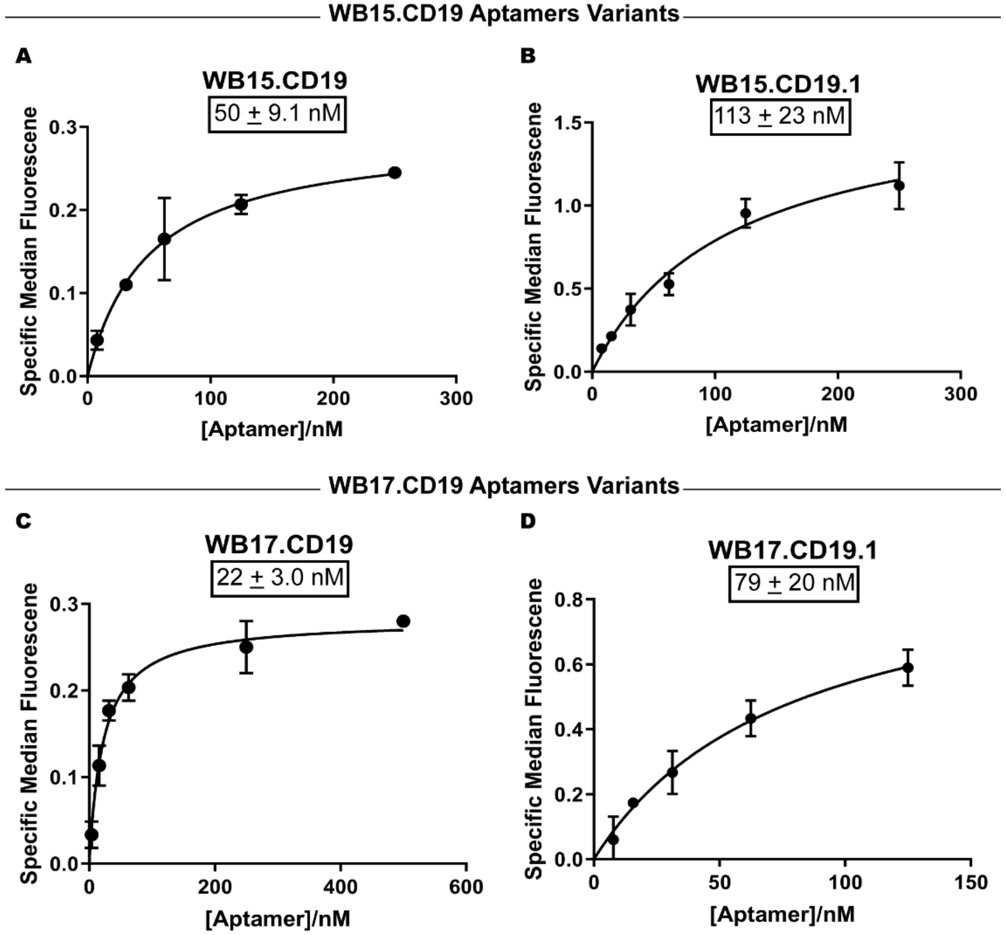


**Figure S2: Apparent Affinities of Parental and Truncated CD19 Aptamers.** Parental CD19 Aptamers and their first truncation variants exhibit unexpected decreases in affinity. (A–D) Binding curves for parental CD19 aptamers WB15.CD19 (A) and WB17.CD19 (C) and their first truncation variants WB15.CD19.1 (B) and WB17.CD19.1 (D). Panel A: WB15.CD19 parental aptamer demonstrated a dissociation constant (Kd) of 50 ± 9.1 nM. Panel B: The truncated variant WB15.CD19.1 exhibited a decreased affinity (Kd = 113 ± 23 nM), highlighting an unexpected reduction in binding efficiency, while panel C shows that the WB17.CD19 parental aptamer had the strongest affinity (Kd = 22 ± 3.0 nM). However, (D) despite truncation leading to decreased affinity, WB17.CD19.1 remained the best candidate among truncated variants with a Kd of 79 ± 20 nM. Notably, WB15.CD19.1 and WB17.CD19.1 differ by only two bases, yet they exhibit distinct binding affinities, emphasizing the importance of nucleotide composition in aptamer performance. Key Insights: Truncation did not improve binding affinity as anticipated, which was unexpected based on prior optimizations. WB17.CD19.1 was identified as the most promising truncated variant despite the decrease in affinity compared to its parent aptamer. Dissociation constants (Kd) and apparent affinity values were determined by plotting the aptamer's specific median fluorescence against its nanomolar (nM) concentration using GraphPad Prism software. The values were calculated using nonlinear regression (curve fitting) using with a one-site specific binding model. Specific median fluorescence was calculated using the formula: $\text{Specific median fluorescence} = \text{Aptamer Median Fluorescence} - \text{Random DNA Median Fluorescence}$. Apparent affinity values were generated using the equation $Y = \frac{(B_{max} \times X)}{(K_d + X)}$, where X represents the aptamer concentration and Y represents the specific median fluorescence. Data are represented as mean ± standard deviation from three independent experiments.

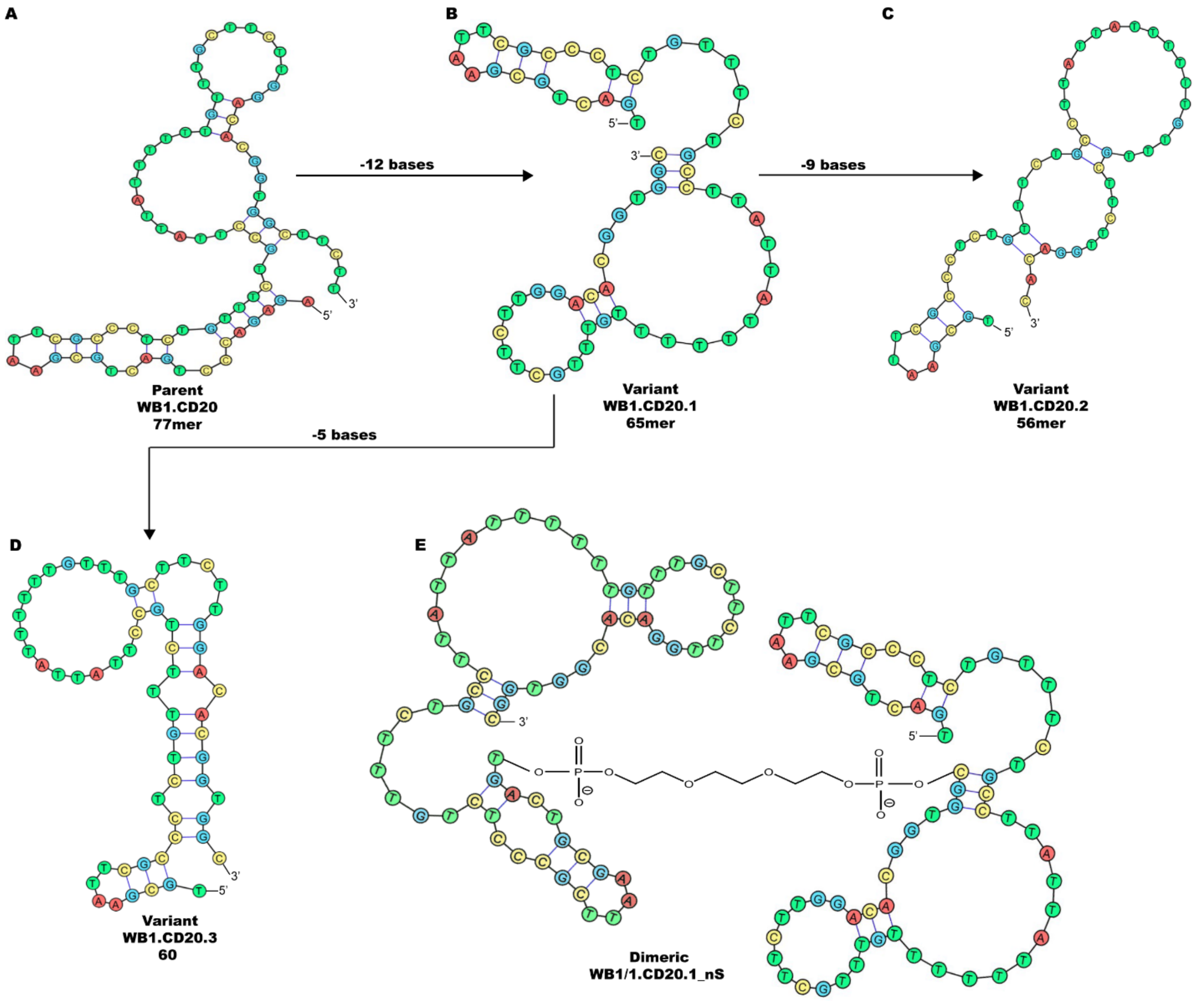


**Figure S3: Predicted Structures and Dimerization of CD20 Aptamers.** Predicted secondary structures of WB1.CD20 aptamer and its truncation variants leading to the engineering of a dimeric aptamer. (A–D) Structural predictions obtained for the parental aptamer WB1.CD20 and its truncated variants. Predictions were generated using m-fold software, NUPACK, visualized with Ribosketch. (A) The parental WB1.CD20 aptamer (77mer) displays a multi-loop structure with regions of high stability. (B) The first truncation variant, WB1.CD20.1 (65mer), maintains the essential structural features of the parent, but eliminates 12 bases to simplify the design, while preserving affinity. (C) The second truncated variant, WB1.CD20.2 (56mer), removes an additional 9 bases, resulting in a compact structure, but with reduced binding affinity. (D) WB1.CD20.3 (60mer) features a unique hairpin-loop structure with truncation optimized for stability and functional performance, but its binding affinity is dramatically decreased. (E) The dimeric aptamer WB1/1.CD20.1_nS was engineered using the best-performing monovalent truncation, WB1.CD20.1. The dimeric design incorporates a 9-PEG linker to connect two monovalent units, enabling bivalent binding to CD20. The engineered dimer retains the individual structural integrity of both monovalent units, facilitating cooperative binding for enhanced affinity and stability. Key Insights: Truncation simplifies the aptamer's structure, while retaining key secondary motifs critical for binding. The dimeric aptamer design leverages the optimized monovalent variant WB1.CD20.1, significantly improving target interaction through multivalent binding. Data represent a stepwise progression from parent to engineered dimeric aptamer.

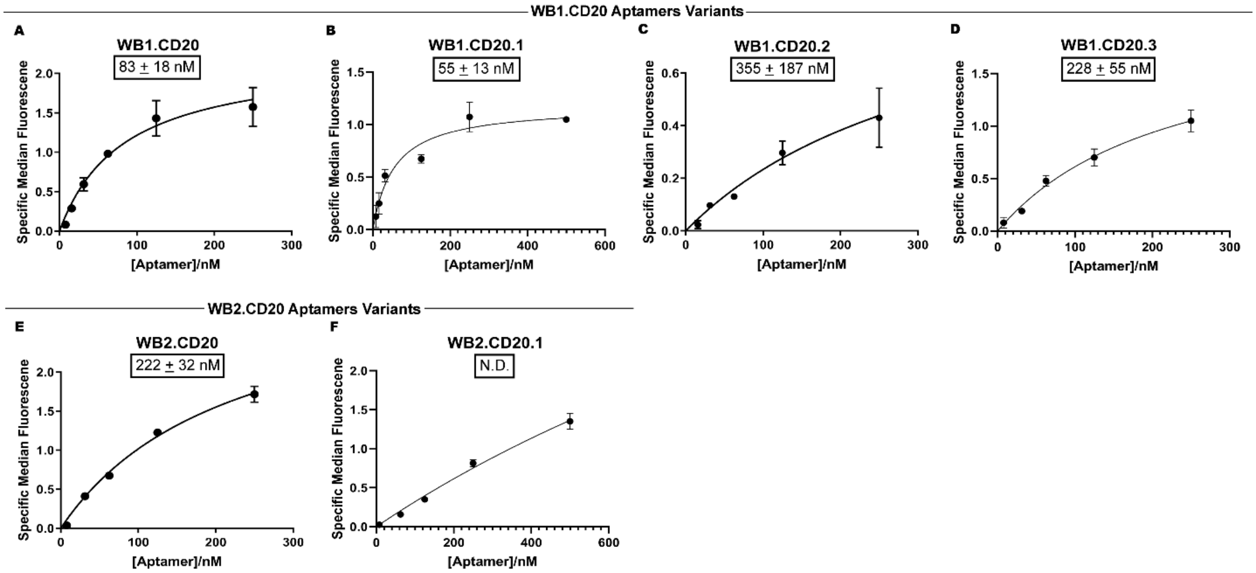


**Figure S4: Apparent Affinities of Parental CD20 Aptamers and their Truncated Variants.** Binding affinities of parental CD20 aptamers WB1.CD20 and WB2.CD20 and the effect of truncation on the affinity. (A–D) Binding curves for WB1.CD20 parental aptamer and its truncated variants (WB1.CD20.1, WB1.CD20.2, and WB1.CD20.3). Truncation of WB1.CD20 improved binding affinity, with WB1.CD20.1 exhibiting the highest affinity (Kd = 55 ± 13 nM), followed by WB1.CD20.3 (Kd = 228 ± 55 nM). WB1.CD20.2 had reduced affinity (Kd = 355 ± 187 nM), highlighting the importance of structural integrity. (E–F) Binding curves for parental aptamer WB2.CD20 and its truncated variant WB2.CD20.1. While WB2.CD20 demonstrated weak affinity (Kd = 222 ± 32 nM), its truncation to WB2.CD20.1 resulted in no detectable binding affinity (Kd = N.D., not determined), highlighting that truncated variant WB1.CD20.1 is the monovalent hit candidate for our anti-CD20 aptamer. Key Findings: Truncation of WB1.CD20 resulted in improved binding affinities for its variants, particularly WB1.CD20.1, compared to the parental aptamer. In contrast, truncation of WB2.CD20 (WB2.CD20.1) resulted in binding affinity that was not determined (N.D.), emphasizing the importance of retaining critical structural regions for optimal target interaction. Dissociation constants (Kd) and apparent affinity values were determined by plotting the aptamer's specific median fluorescence against its nanomolar (nM) concentration using GraphPad Prism software. The values were calculated using nonlinear regression (curve fitting) using with a one-site specific binding model. Specific median fluorescence was calculated using the formula: $\text{Specific median fluorescence} = \text{Aptamer Median Fluorescence} - \text{Random DNA Median Fluorescence}$. Apparent affinity values were generated using the equation $Y = \frac{(B_{max} \times X)}{(K_d + X)}$, where X represents the aptamer concentration and Y represents the specific median fluorescence. Data are represented as mean ± standard deviation from three independent experiments. N.D. indicates values that were not determined.

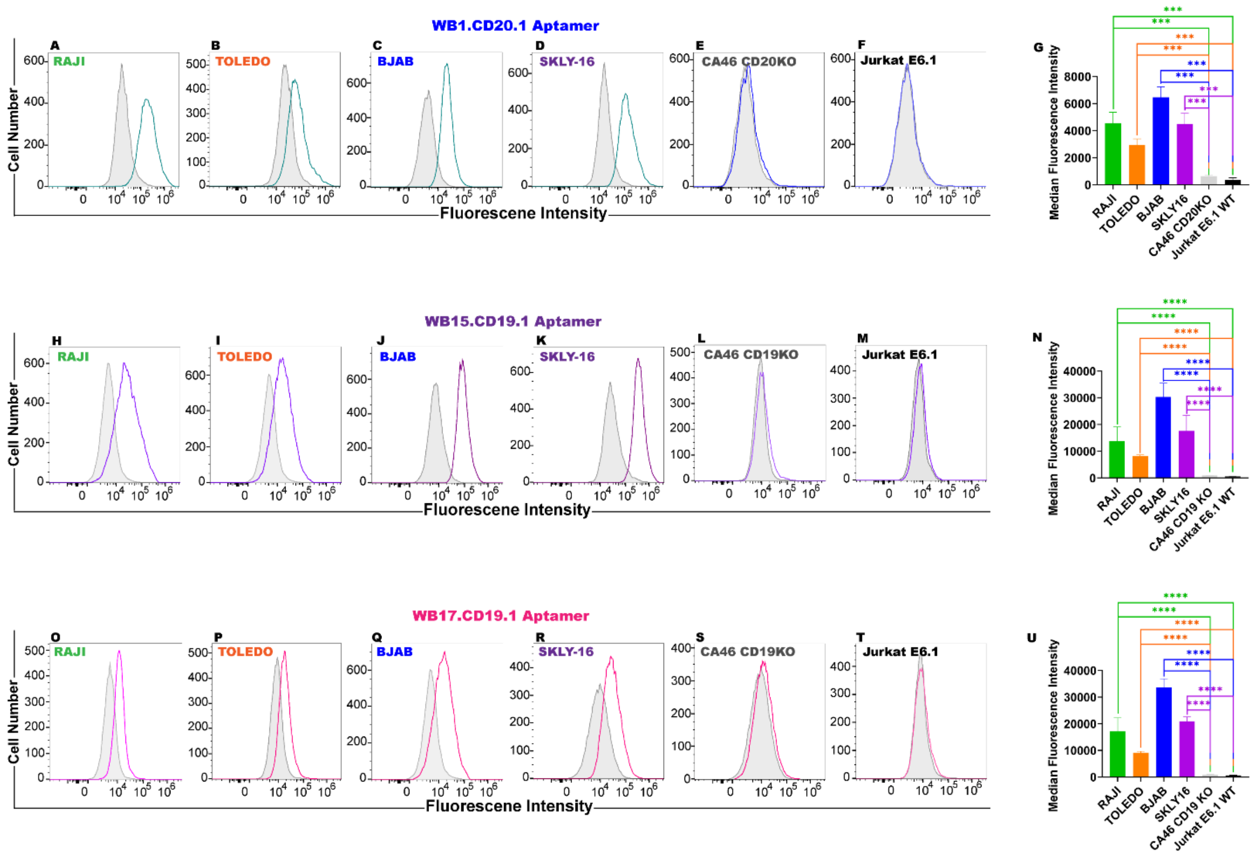


**Figure S5: Binding analysis of Truncated Monovalent Aptamers to CD19 and CD20.** Binding of truncated monovalent WB1.CD20.1, WB15.CD19.1, and WB17.CD19.1 aptamers with various cell lines. (A–G) Binding analysis of WB1.CD20.1 aptamer (Blue): (A–F) Fluorescence intensity histograms for CD20-positive cell lines (RAJI (A), TOLEDO (B), BJAB (C), SKLY-16 (D)) compared to knockdown (CA46 CD20KO (E)) and negative (Jurkat E6.1 (F)) cell lines. WB1.CD20.1 demonstrates strong binding to CD20-positive cell lines, but negligible binding to CD20KO and negative controls. (G) Bar graph of median fluorescence intensities highlight the specificity of WB1.CD20.1 for CD20-positive cells and its significantly reduced binding to knockdown and negative lines. (H–N) Binding analysis of WB15.CD19.1 aptamer (Purple): (H–M) Fluorescence intensity histograms for CD19-positive cell lines (RAJI (H), TOLEDO (I), BJAB (J), SKLY-16 (K)) compared to knockdown (CA46 CD19KO (L)) and negative (Jurkat E6.1 (M)) cell lines. WB15.CD19.1 shows specific binding to CD19-positive cells and minimal binding to CD19KO or negative controls. (N) Bar graph of median fluorescence intensities confirms strong binding specificity of WB15.CD19.1 to CD19-positive cells. (O–U) Binding analysis of WB17.CD19.1 aptamer (Pink): (O–T) Fluorescence intensity histograms for CD19-positive cell lines (RAJI (O), TOLEDO (P), BJAB (Q), SKLY-16 (R)) compared to knockdown (CA46 CD19KO (S)) and negative (Jurkat E6.1 (T)) cell lines. WB17.CD19.1 exhibits robust binding to CD19-positive cells, but only minimal interaction with CD19KO or negative controls. (U) Bar graph of median fluorescence intensities further supports the strong specificity of WB17.CD19.1 for CD19-positive cell lines. Key Insights: All three truncated monovalent aptamers (WB1.CD20.1, WB15.CD19.1, and WB17.CD19.1) demonstrate high specificity for their respective targets (CD20 or CD19) in positive cell lines. Binding to knockdown (KO) and negative control cell lines was significantly reduced, confirming specificity. Median fluorescence intensity was calculated using the formula: $\text{Median fluorescence intensity} = \text{Aptamer Median Fluorescence} - \text{Random DNA Median Fluorescence}$. The aptamer and random median fluorescence values corresponds to the median fluorescence observed in their respective histograms. Bar graphs represent median ± standard deviation from three independent experiments with statistical significance indicated (****p < 0.0001).

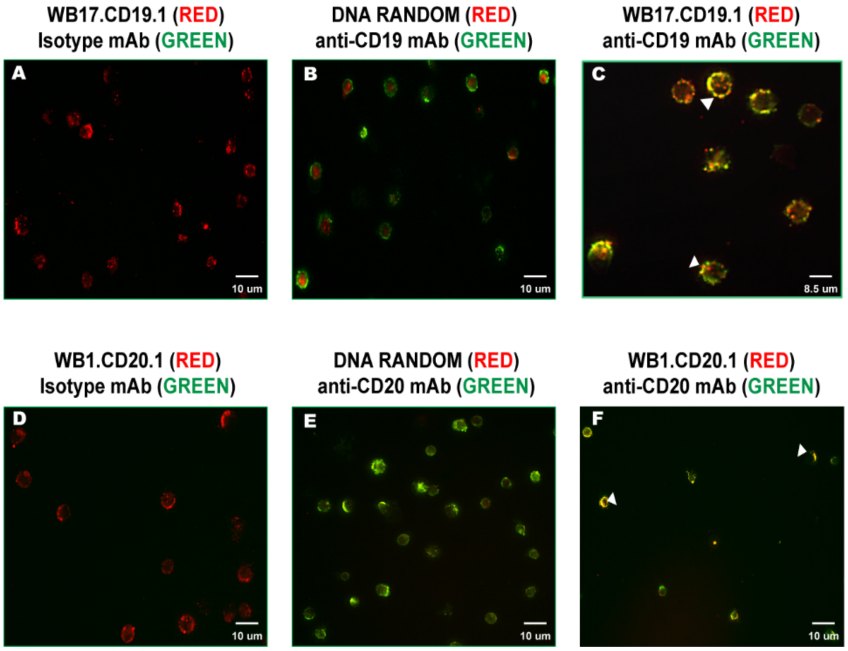


**Figure S6: Colocalization of Truncated Monovalent Aptamers with Anti-CD19 and Anti-CD20 Antibodies**. Colocalization of anti-CD19 and anti-CD20 antibodies with truncated monovalent aptamers WB17.CD19.1 and WB1.CD20.1 at the cell surface of Toledo cells. (A–C) Colocalization of anti-CD19 antibody (Green) with WB17.CD19.1 aptamer (Red): (A) Binding of WB17.CD19.1 aptamer visualized in red. (B) Binding of Anti-CD19 antibody visualized in green. (C) Merged image showing colocalization of WB17.CD19.1 aptamer with anti-CD19 antibody at the cell surface membrane, as indicated by yellow regions (arrowheads), confirming their spatial overlap. (D–F) Colocalization of anti-CD20 antibody (Green) with WB1.CD20.1 aptamer (Red): (D) Binding of WB1.CD20.1 aptamer visualized in red. (E) Binding of Anti-CD20 antibody visualized in green. (F) Merged image showing colocalization of WB1.CD20.1 aptamer with anti-CD20 antibody at the cell surface membrane, as indicated by yellow regions (arrowheads), confirming their spatial overlap. Key Insights: Both truncated monovalent aptamers, WB17.CD19.1 and WB1.CD20.1, colocalize with their respective antibodies, validating their binding specificity to CD19 and CD20 on the surface membrane of Toledo cells. Colocalization was visualized using a Nikon TiE inverted microscope, indicating that the aptamer and antibody target the same surface structures. Images were generated using fixed Toledo cells stained with truncated monovalent aptamers and respective anti-CD19 or anti-CD20 antibodies. Images were acquired using a Nikon TiE inverted microscope (Nikon Inc., Melville, NY). Colocalization was visualized by the overlap of red (aptamer) and green (antibody) signals, resulting in yellow fluorescence. Scale Bars: 8.5 and 10 μm.

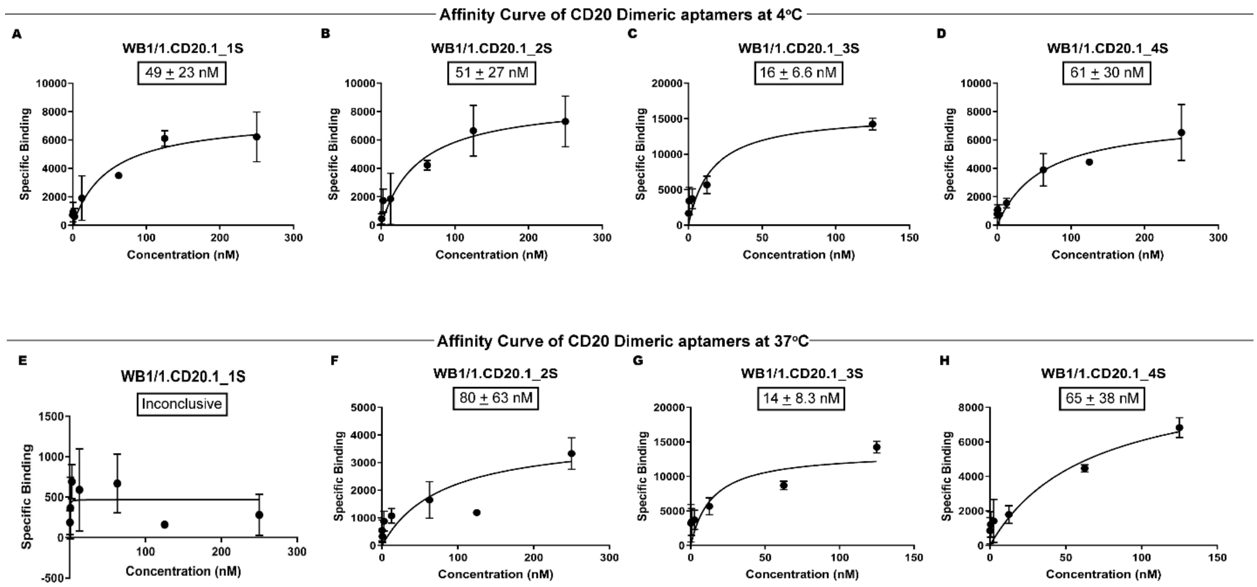


**Figure S7: Apparent Affinities of Homodimeric CD20 Aptamers with Variable Spacers.** Affinity of homodimeric CD20 aptamers with variable spacers highlighting optimal spacer configuration (3 Spacer, SP9) for target interaction. (A–D) Affinity curves for homodimeric WB1/1.CD20.1 variants measured at 4°C: (A) WB1/1.CD20.1_1S: A single-spacer configuration showed reduced binding affinity with a dissociation constant (Kd) of 49 ± 23 nM. (B) WB1/1.CD20.1_2S: Two-spacer configuration with a Kd of 51 ± 27 nM, showing moderate binding efficiency. (C) WB1/1.CD20.1_3S (SP9): The best-performing variant with a 9-PEG spacer providing an optimal distance of 3.96 nm between monomer units, achieving the highest binding affinity (Kd = 16 ± 6.6 nM). (D) WB1/1.CD20.1_4S: Four spacers showed weaker binding compared to the 3-spacer variant with a Kd of 61 ± 30 nM. (E–H) Affinity curves for the same WB1/1.CD20.1 variants measured at 37°C: (E) WB1/1.CD20.1_1S exhibited inconclusive binding at physiological temperature, indicating an inability to determine performance under these conditions. (F) WB1/1.CD20.1_2S with moderate binding at 37°C (Kd = 80 ± 63 nM). (G) WB1/1.CD20.1_3S (SP9) maintained strong binding at 37°C with a Kd of 14 ± 8.3 nM, confirming its superior stability and binding performance across temperatures. (H) WB1/1.CD20.1_4S displayed weaker affinity at 37°C (Kd = 65 ± 38 nM). Key Insights: The WB1/1.CD20.1_3S variant (SP9) with a 9-PEG spacer providing a 3.96 nm distance between the two truncated monovalent aptamers demonstrates the best binding affinity at both 4°C and 37°C. Spacer configuration and length are critical for optimizing multivalent interactions, with shorter or excessive spacers leading to suboptimal binding. Binding assays were conducted using flow cytometry. Incubation was performed at 4°C and 37°C. Specific binding was calculated using the formula: $\text{Specific Mean fluorescence} = \text{Aptamer Mean Fluorescence} - \text{Random DNA Mean Fluorescence}$. The aptamer and random mean fluorescence values correspond to the mean fluorescence of the respective histograms. Apparent affinity values were generated by plotting the specific aptamer binding against the aptamer concentration using GraphPad Prism software with a nonlinear fit and one-site specifici binding model: $Y = \frac{(B_{max} \times X)}{(K_d + X)}$, where X represents the aptamer concentration and Y represents the specific mean fluorescence. Dissociation constants (Kd) were determined using nonlinear regression (curve-fitting) analysis. N.D. indicates values that were not determined. Data are represented mean ± standard deviation from three independent experiments.

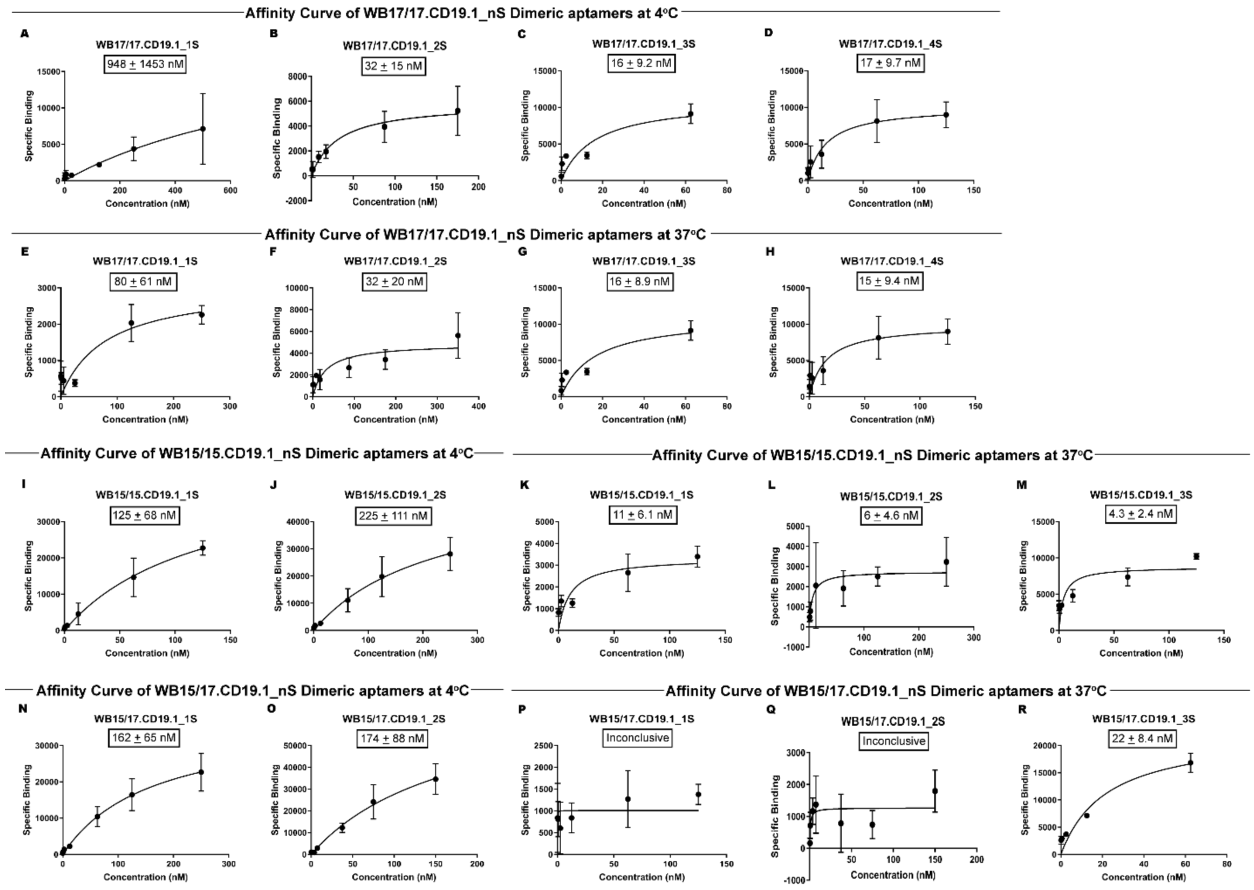


**Figure S8: Apparent Affinities of Homodimeric and Heterodimeric CD19 Aptamers.** Affinity of homodimeric and heterodimeric CD19 aptamers across spacer configurations at 4°C and 37°C. (A–D) Affinity curves for WB17/17.CD19.1 homodimeric aptamers at 4°C: (A) WB17/17.CD19.1_1S: A single- spacer configuration exhibited poor binding (Kd = 948 ± 1453 nM). (B) WB17/17.CD19.1_2S: Two-spacer configuration improved binding (Kd = 32 ± 15 nM). (C) WB17/17.CD19.1_3S (SP9): The optimal configuration achieved the highest affinity (Kd = 16 ± 9.2 nM), with a 9-PEG spacer length of 3.96 nm. (D) WB17/17.CD19.1_4S: Four-spacer configuration demonstrated weaker binding (Kd = 17 ± 9.2 nM). (E–H) Affinity curves for WB17/17.CD19.1 homodimeric aptamers at 37°C: (E) WB17/17.CD19.1_1S: Reduced binding efficiency at physiological temperature (Kd = 80 ± 61 nM). (F) WB17/17.CD19.1_2S: Enhanced binding with Kd = 32 ± 20 nM. (G) WB17/17.CD19.1_3S (SP9): Maintained strong binding with Kd = 16 ± 8.9 nM, confirming optimal performance. (H) WB17/17.CD19.1_4S: Maintained binding at 37°C (Kd = 15 ± 9.4 nM), but plateaued at 3 SP9 (WB17/17.CD19.1_3S: G) as the hit candidate. (I–M) Affinity curves for WB15/15.CD19.1 homodimeric aptamers at 4°C and 37°C: (I, J) WB15/15.CD19.1_1S and _2S showed reduced binding efficiency with higher Kd values. (K, L) Improved binding performance at 37°C. (M) WB15/15.CD19.1_3S (SP9) displayed the best binding affinity at 37°C (Kd = 4.3 ± 2.4 nM). (N–R) Affinity curves for WB15/17.CD19.1 heterodimeric aptamers at 4°C and 37°C: (N) WB15/17.CD19.1_1S showed weaker binding performance (Kd = 162 ± 65 nM). (O) WB15/17.CD19.1_2S maintained binding (Kd = 174 ± 88 nM). (R) WB15/17.CD19.1_3S (SP9) showed optimal binding with the strongest affinity at 37°C (Kd = 22 ± 8.4 nM). Key Insights: The SP9 configuration (3 spacer, 9-PEG, 3.96 nm) consistently demonstrated the strongest binding for both homodimeric and heterodimeric CD19 aptamers. Spacer length and configuration significantly impact binding efficiency, with shorter or excessive spacers reducing performance. Binding assays were performed using flow cytometry. Incubation at 4°C and 37°C. Specific binding was calculated using the formula: $\text{Specific Mean fluorescence} = \text{Aptamer Mean Fluorescence} - \text{Random DNA Mean Fluorescence}$. The aptamer and random mean fluorescence values correspond to the mean fluorescence of the respective histograms. Apparent affinity values were generated by plotting the specific aptamer binding against the aptamer concentration using GraphPad Prism software with a nonlinear fit and one-site specifici binding model: $Y = \frac{(B_{max} \times X)}{(K_d + X)}$, where X represents the aptamer concentration and Y represents the specific mean fluorescence. Dissociation constants (Kd) were determined using nonlinear regression (curve-fitting) analysis. Data are represented mean ± standard deviation from three independent experiments.

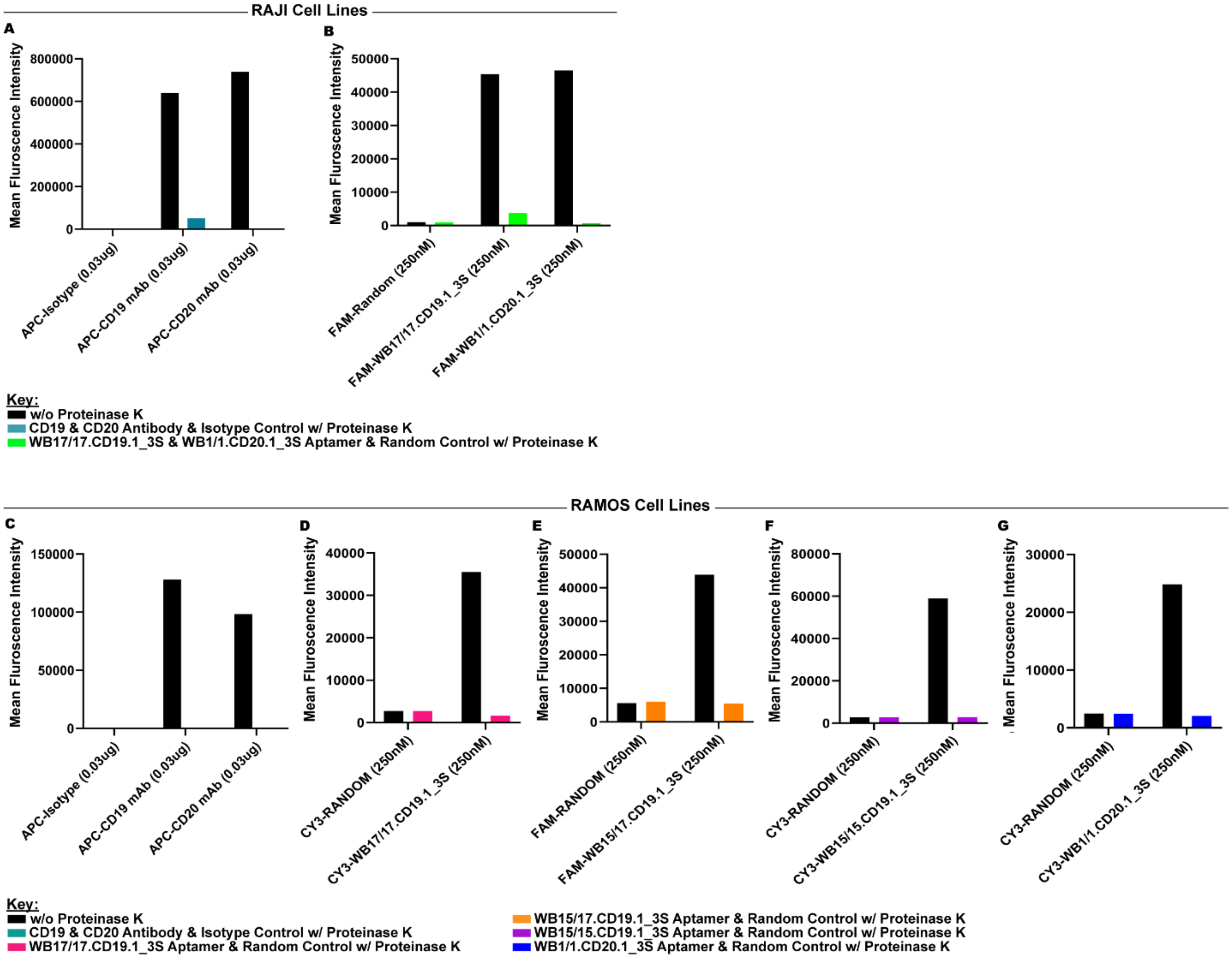


**Figure S9: Proteinase K Treatment Efficiency on CD19 and CD20 in Raji and Ramos Cells.** Proteinase K Treatment on Raji and Ramos Cells demonstrates efficient cleavage of extracellular CD19 and CD20. (A–B) Raji Cells: (A) Mean fluorescence intensity (MFI) of APC-CD19 and APC-CD20 antibodies on untreated and Proteinase K-treated Raji cells. Treatment with 1 mg/mL Proteinase K effectively removed CD19 and CD20 from the surface membrane. (B) MFI of dimeric aptamers (FAM-WB17/17.CD19.1_3S and FAM-WB1/CD20.1_3S) on Raji cells, confirming surface cleavage of both aptamer-bound CD19 and CD20 targets. (C–G) Ramos Cells: (C) MFI of APC-CD19 antibody on untreated and Proteinase K-treated Ramos cells, demonstrating extracellular cleavage of CD19 and CD20. (D–G) MFI of dimeric aptamers (CY3-WB17/17.CD19.1_3S (Pink, D), FAM-WB15/17.CD19.1_3S (Orange, E), CY3-WB15/15.CD19.1_3S (Purple, F), and CY3-WB1/CD20.1_3S (Blue, G)) with random DNA controls (FAM-RANDOM and CY3-RANDOM), confirming effective cleavage of surface-bound CD19 and CD20 at 1 mg/mL Proteinase K concentration. Key Insights: Proteinase K treatment at 1 mg/mL efficiently removed extracellular CD19 and CD20 from Raji and Ramos cells, making them suitable for internalization studies. Proteinase K treatment was applied twice for 10 minutes after antibody or aptamer binding, ensuring robust removal of extracellular targets. Cells were stained with antibodies or bound with aptamers, treated with Proteinase K (1 mg/mL), and analyzed by flow cytometry to measure MFI. Data represent mean ± standard deviation from three independent experiments.

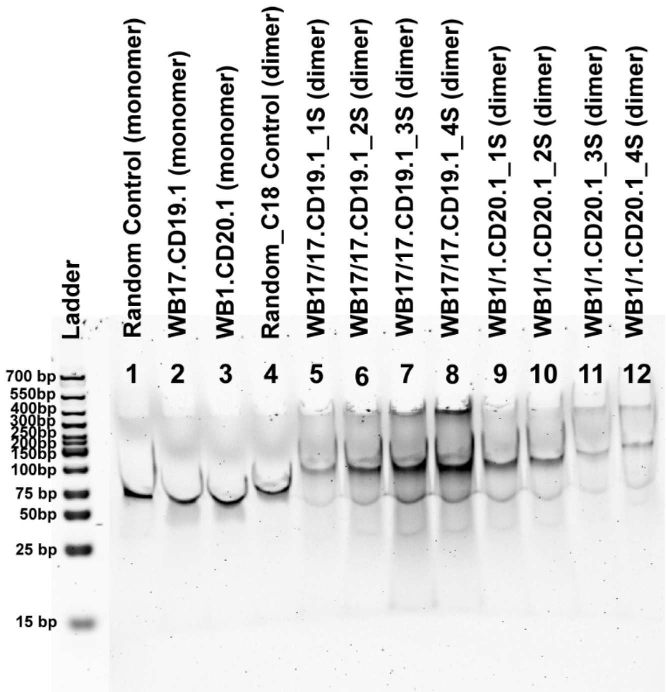


7.5% Denaturing PAGES

**Figure S10: Serum Stability of Monovalent and Dimeric Aptamers.** This figure demonstrates the stability of monomeric and dimeric aptamers after 24 hours of incubation in 10% fetal bovine serum (FBS) media at 37°C. Aptamers, including WB17.CD19.1 (monomer), WB1.CD20.1 (monomer), and various dimeric variants (e.g., WB17/17.CD19.1_1S, WB17/17.CD19.1_2S, WB17/17.CD19.1_3S, WB17/17.CD19.1_4S, WB1/1.CD20.1_1S, WB1/1.CD20.1_2S, WB1/1.CD20.1_3S, and WB1/1.CD20.1_4S), were analyzed using a 7.5% denaturing polyacrylamide gel. The gel showed that none of the tested aptamers displayed any visible degradation bands, confirming their structural integrity and resistance to serum nucleases. A random control (monomer and dimer) was included for comparison. Aptamers were incubated with 20% FBS media (1:1 ratio), followed by analysis on a 7.5% denaturing PAGE gel. Electrophoresis revealed sharp, intact bands for all aptamers, indicating that both monomeric and dimeric forms remain stable under these conditions. The absence of smearing or additional bands suggests that aptamers are resistant to degradation by serum proteins, making them robust candidates for both *in vitro* and potential *in vivo* applications. This result underscores the high stability and functional viability of these aptamers when exposed to physiological conditions.

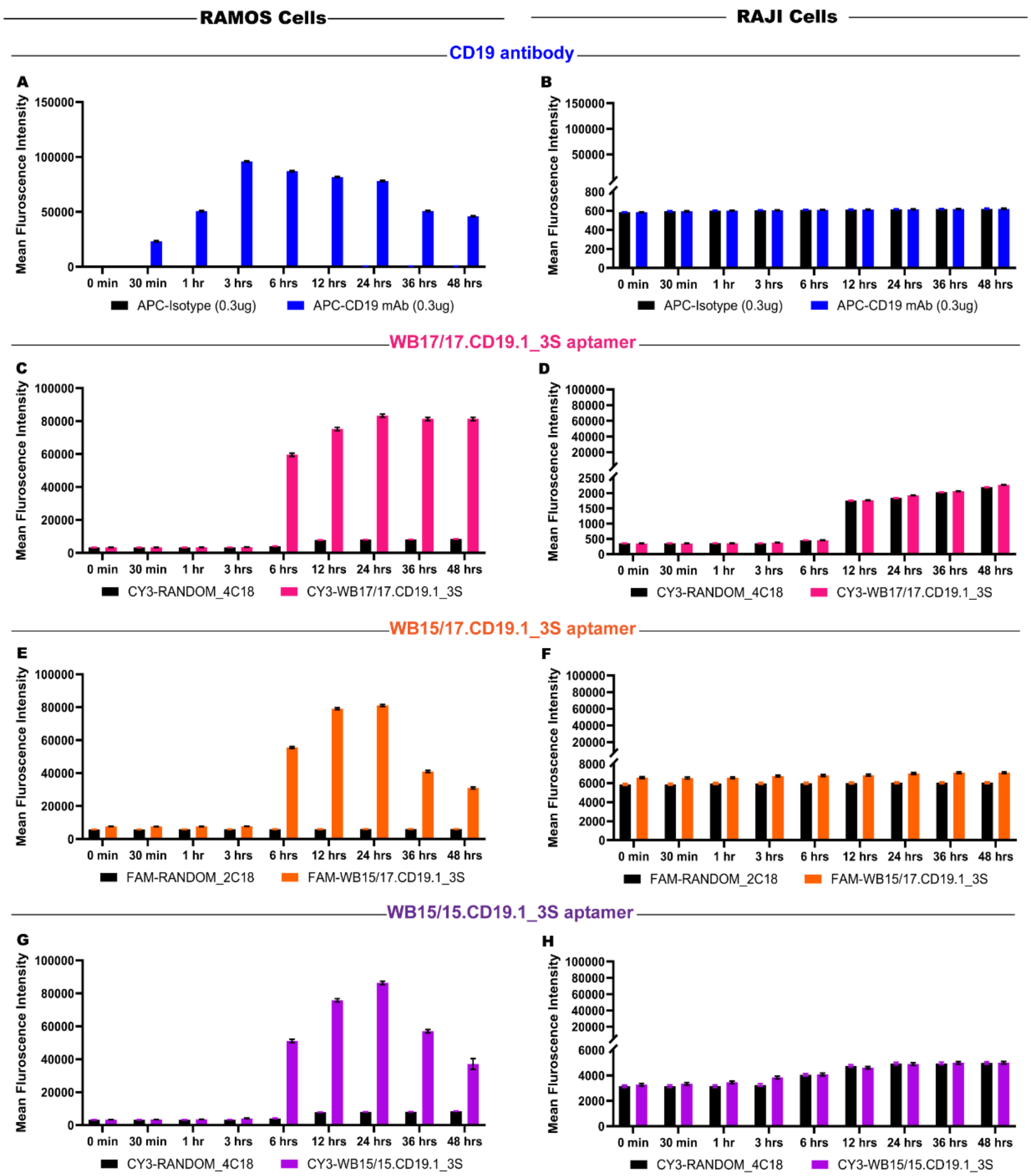


**Figure S11: Fluorescent Labeling Does Not Influence Internalization of CD19 Antibody and Aptamers in Ramos and Raji Cells.** (A-B) CD19 Antibody Internalization: (A) In Ramos cells, APC-CD19 antibody demonstrates robust internalization over 48 hours compared to the APC-labeled isotype control, confirming efficient CD19-specific internalization. The rapid uptake is complete after 3 hours. (B) In Raji cells, no significant internalization of APC-CD19 mAb is observed compared to the APC-labeled isotype control. These results confirm that the fluorescent label (APC) does not influence internalization. (C-H) Aptamer Internalization: (C, E, G) In Ramos cells, dimeric CD19 aptamers (WB17/17.CD19.1_3S (pink, C), WB15/17.CD19.1_3S (orange, E), and WB15/15.CD19.1_3S (purple, G)) labeled with either Cy3 or FAM show time-dependent internalization over 48 hours, with significantly higher fluorescence intensities compared to random DNA controls. Complete uptake occurs in 24 hours. (D, F, H) In Raji cells, dimeric aptamers show no internalization, with fluorescence intensities comparable to those of random DNA controls. Importantly, random DNA controls (Cy3- or FAM-labeled) do not internalize in either Ramos or Raji cells, further confirming that the fluorescent label does not affect internalization. This figure highlights that the choice of fluorescent label (APC, Cy3, or FAM) does not influence internalization. Specific internalization is driven by CD19-targeting aptamers or antibodies, as demonstrated by comparisons with isotype controls and random DNA aptamers. Internalization is efficient in Ramos cells, while no internalization occurs in Raji cells, regardless of presents of the fluorescent label. Controls, including isotype antibodies and random DNA aptamers, validate the specificity and lack of fluorescent label involvement in observed internalization. Ramos and Raji cells were stained with APC-CD19 antibody or incubated with dimeric CD19 aptamers (WB17/17.CD19.1_3S, WB15/17.CD19.1_3S, and WB15/15.CD19.1_3S) and analyzed at eight time points. Cells were treated with Proteinase K to remove extracellular CD19, ensuring that only internalized molecules were detected. Data are presented as mean fluorescence intensities ± standard deviation from three independent experiments.

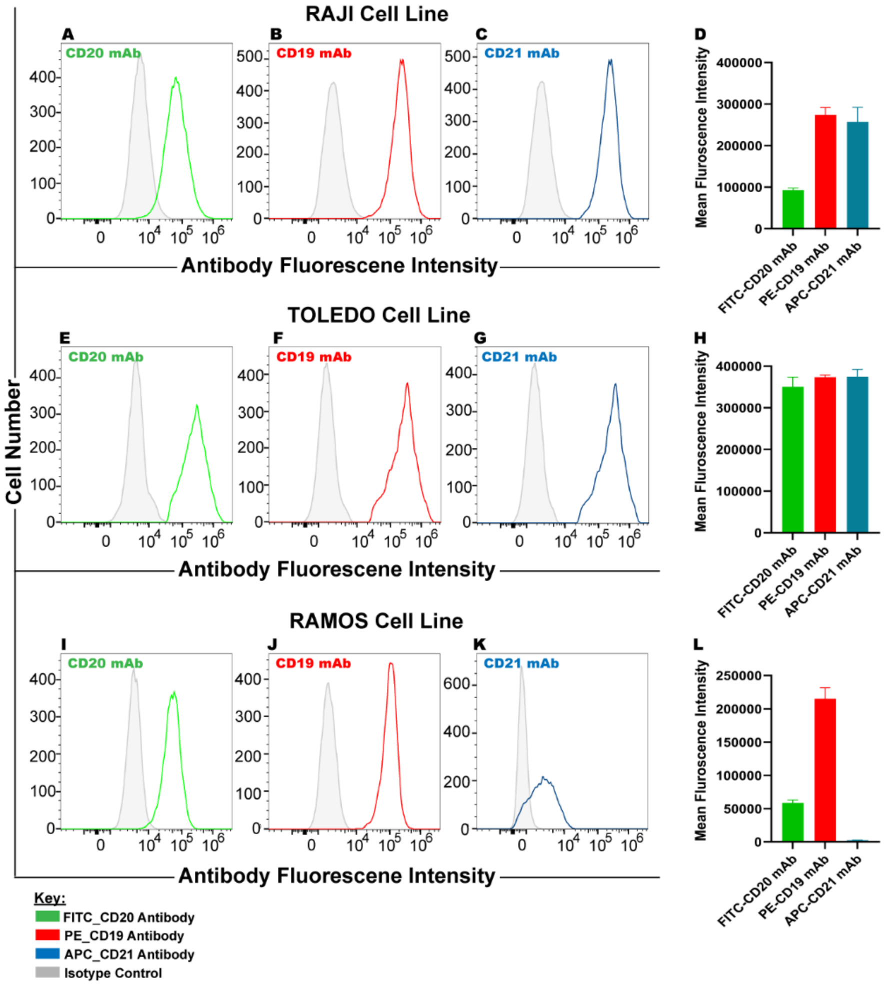


**Figure S12: Analysis of CD19 and CD21 Expression Across Raji, Toledo, and Ramos Cells**. The analysis of CD19 and CD21 expression across Raji, Toledo, and Ramos cells using flow cytometry to Identify the best model for CD19 internalization studies. (A–D) Antibody staining in Raji cells: (A–C) Fluorescence intensity histograms for CD20 (Green), CD19 (Red), and CD21 (Light Blue) antibodies, showing strong expression of all three markers, including CD21, the co-receptor of CD19. (D) Bar graph of mean fluorescence intensities confirms the expression of CD21 in Raji cells. The presence of CD21 suggests interference with CD19 internalization, making Raji cells unsuitable for studying CD19-specific internalization. (E–H) Antibody staining in Toledo cells: (E–G) Fluorescence intensity histograms for CD20 (Green), CD19 (Red), and CD21 (Light Blue) antibodies, indicating high expression of all markers, including CD21. (H) Bar graph of mean fluorescence intensities highlights the expression of CD21, which also makes Toledo cells unsuitable for CD19-specific internalization studies owing to the role of CD21 in influencing CD19 dynamics. (I–L) Antibody staining in Ramos cells: (I–K) Fluorescence intensity histograms for CD20 (Green), CD19 (Red), and CD21 (Light Blue) antibodies. Ramos cells exhibit strong CD19 expression, but lack CD21 expression. (L) Bar graph of mean fluorescence intensities confirms the absence of CD21 in Ramos cells, identifying Ramos as the optimal model for studying CD19-specific internalization without interference from CD21. Key Insights: Raji and Toledo cells, both positive for CD21, are unsuitable models for CD19-specific internalization studies because CD21 inhibits CD19 internalization dynamics. Ramos cells, which lack CD21 expression, provide a clean system to study CD19-specific internalization without confounding effects from the co-receptor. Cells were stained with fluorescently labeled antibodies (FITC-CD20, PE-CD19, and APC-CD21), and fluorescence intensities were measured using flow cytometry to analyze binding events. Mean fluorescence intensities (MFI) were calculated using the formula: $\mathrm{MFI} = \mathrm{Antibody\ Mean\ Fluorescence} - \mathrm{Isotype\ Control\ Mean\ Fluorescence}$, where the antibody and isotype control mean fluorescence values correspond to the mean fluorescence observed in their respective histograms. MFIs were plotted to compare marker expression across cell lines. Error bar represents mean ± standard deviation from three independent experiments.

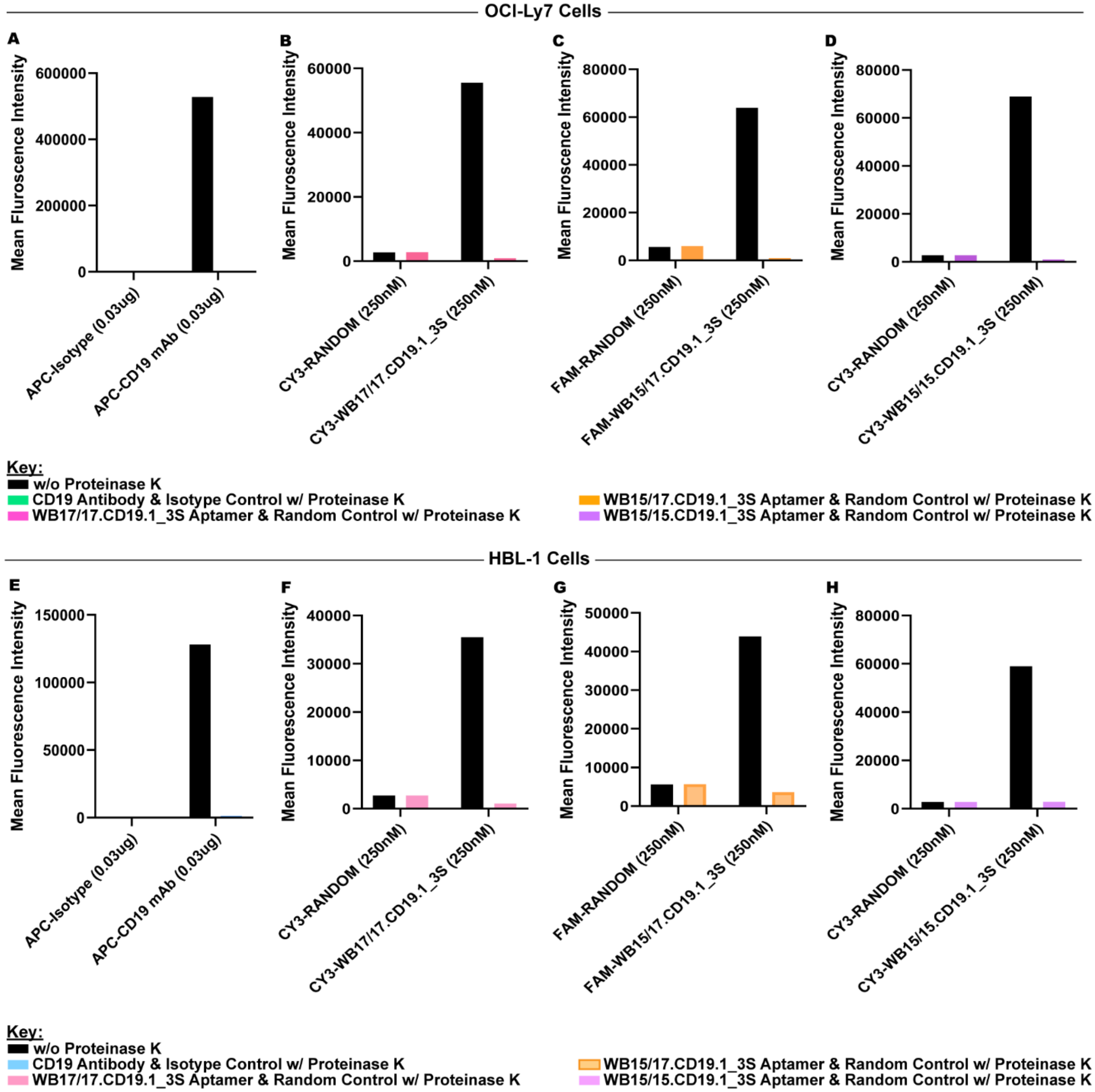


**Figure S13: Proteinase K Treatment on OCI-Ly7 and HBL-1 Cells.** Proteinase K Treatment on OCI-Ly7 and HBL-1 Cells Demonstrates Efficient Cleavage of Extracellular CD19. (A–D) OCI-Ly7 Cells: (A) MFI of APC-CD19 antibody and Isotype control on untreated and Proteinase K-treated OCI-Ly7 cells. A higher Proteinase K concentration of 3 mg/mL was required to achieve complete cleavage of extracellular CD19. (B-D) MFI of dimeric aptamers (CY3-WB17/17.CD19.1_3S (Pink, B), FAM-WB15/17.CD19.1_3S (Orange, C), and CY3-WB15/15.CD19.1_3S (Purple, D)) with random DNA controls (FAM-RANDOM and CY3-RANDOM), confirming successful cleavage of surface-bound targets at a concentration of 3 mg/mL Proteinase K. (E-H) HBL-1 Cells: (E) MFI of APC-CD19 antibody and isotype control on untreated and Proteinase K-treated HBL-1 cells. A Proteinase K concentration of 3 mg/mL was required to achieve complete cleavage of extracellular CD19. (F-H) MFI of dimeric aptamers (CY3-WB17/17.CD19.1_3S (Light Pink, F), FAM-WB15/17.CD19.1_3S (Light Orange, G), and CY3-WB15/15.CD19.1_3S (Light Purple, H)) with random DNA controls (FAM-RANDOM and CY3-RANDOM), confirming successful cleavage of surface-bound targets at a concentration of 3 mg/mL Proteinase K. Key Insights: Proteinase K treatment at a concentration of 3 mg/mL results in complete cleavage of the surface of OCI-Ly7 and HBL-1 cells, both diffuse large B-cell lymphoma (DLBCL). Either potentially higher protein abundance or differences in membrane composition required a much higher Proteinase K concentration. Proteinase K treatment was applied twice for 10 minutes after antibody or aptamer binding, ensuring robust removal of extracellular targets, making them suitable for internalization studies. Cells were stained with antibodies or bound with aptamers, treated with Proteinase K (3 mg/mL), and analyzed by flow cytometry to measure MFI. Data represent mean ± standard deviation from three independent experiments.

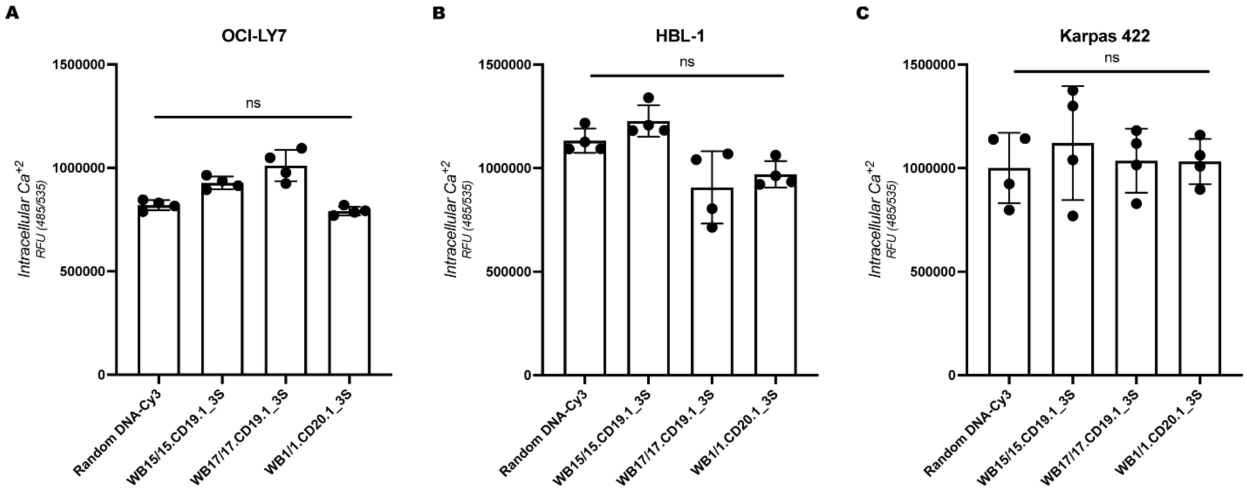


**Figure S14: Intracellular Calcium Release Analysis in DLBCL Cell Lines.** Intracellular calcium release measured in three cell lines, OCI-LY7 (A), HBL-1 (B), and Karpas 422 (C), treated with random DNA control, WB15/15.CD19.1_3S, WB17/17.CD19.1_3S, and WB1/1.CD20.1_3S. Cells were preloaded with the calcium-sensitive dye Fluo-4 AM, and fluorescence emission at 535 nm was measured following aptamer binding. No significant (ns) changes in intracellular calcium release were observed among treatments across all cell lines. Error bars represent the standard deviation (SD) of three independent experiments, each performed in triplicate. Statistical analysis was performed using one-way ANOVA.

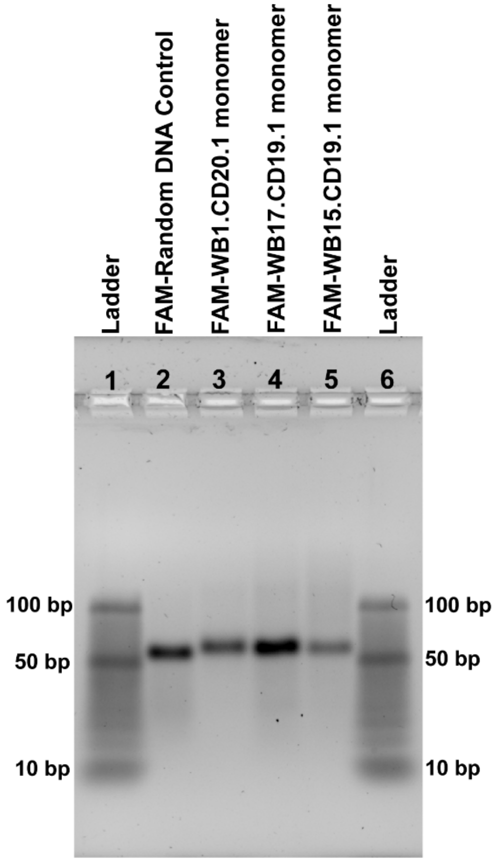


**Figure S15: Analysis of Monovalent CD19 and CD20 Aptamers after HPLC purification.** This figure demonstrates the electrophoretic mobility and purity of the monovalent aptamers WB1.CD20.1 (well 3), WB17.CD19.1 (well 4), and WB15.CD19.1 (well 5) alongside FAM-labeled random DNA control (well 1). Samples were resolved with a 3.5% agarose gel. Lane 2: FAM-Random DNA Control, synthesized and obtained directly from Integrated DNA Technologies (IDT), showing expected mobility for a single-stranded DNA fragment. Lane 3: FAM-WB1.CD20.1 monovalent aptamer, purified via HPLC, displaying a single sharp band indicating high purity. Lane 4: FAM-WB17.CD19.1 monovalent aptamer, purified via HPLC, with a single band demonstrating purity and expected fragment size. Lane 5: FAM-WB15.CD19.1 monovalent aptamer, purified via HPLC, showing a single band corresponding to the expected size. Lane 1 and 6: DNA ladder for reference, marking the 10 bp, 50 bp, and 100 bp bands. Samples were prepared at a concentration of 200 nM. Gel electrophoresis was performed on a 3.5% agarose gel at 120 V for 45 minutes. Post-staining was performed with SYBR Gold for 40 minutes to enhance band visualization. Key Insights: HPLC purification of the monovalent aptamers WB1.CD20.1, WB17.CD19.1, and WB15.CD19.1 resulted in high-purity products, as indicated by the presence of single sharp bands. The random DNA control obtained from IDT also exhibits expected purity and size. Uniformity and sharpness of the bands validate the high quality of the synthesized and purified aptamers, ensuring reliability for applications.

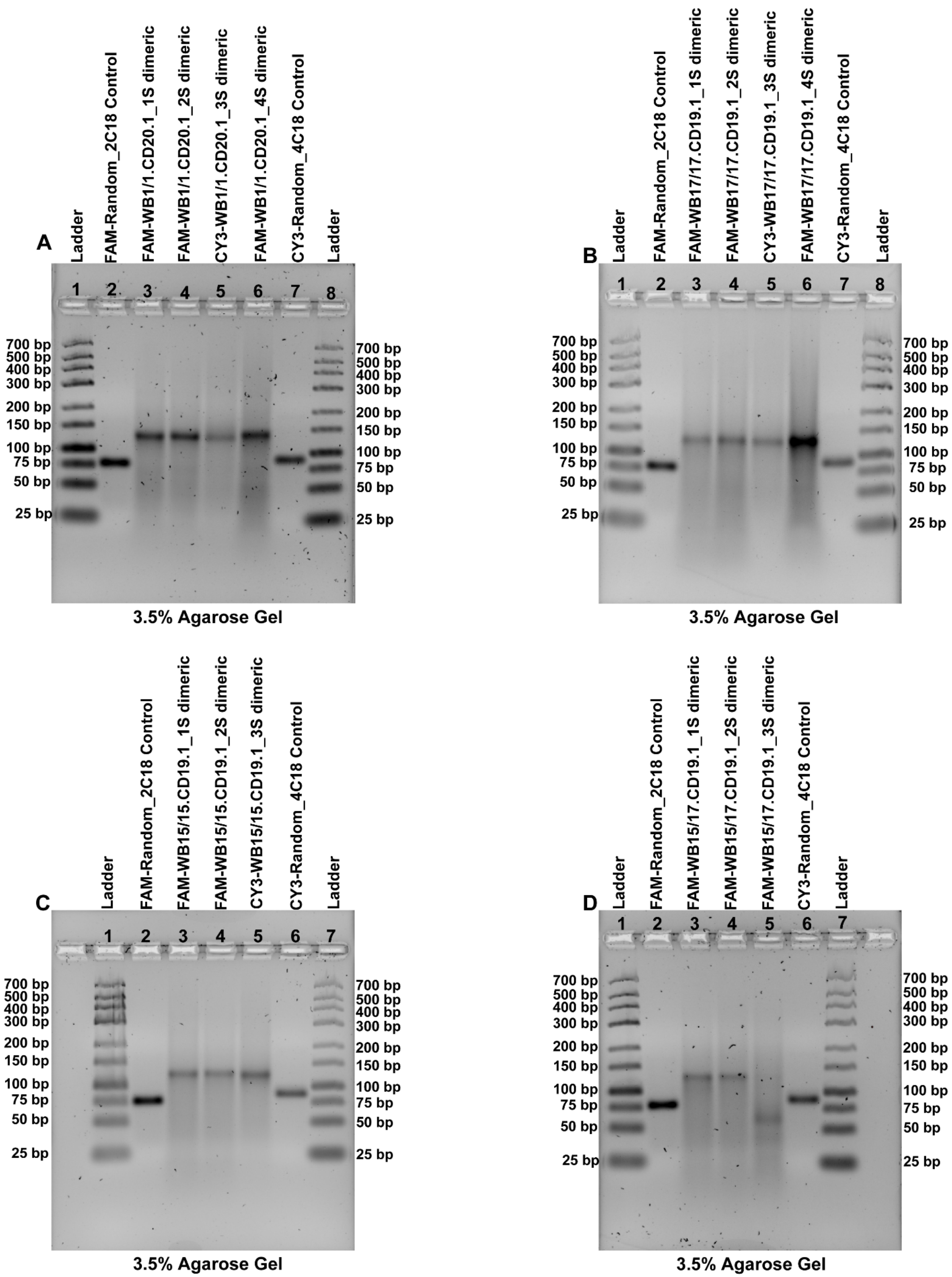


**Figure S16: Analysis of Dimeric CD19 and CD20 Aptamers Following HPLC Purification Compared to Random DNA Controls.** (A–D) 3.5% agarose gel showing the electrophoretic mobility and purity of FAM-labeled and CY3-labeled dimeric aptamers alongside random DNA controls. (A) WB1/1.CD20.1 Dimeric Aptamers: Lane 2: FAM-

Random DNA Control (2C18). Lanes 3–6: Dimeric variants of WB1/1.CD20.1_nS with varying spacer lengths (1S, 2S, 3S, 4S) demonstrate sharp single bands corresponding to their expected sizes, indicating high purity after HPLC purification. Lane 7: CY3-Random DNA Control (4C18). (B) WB17/17.CD19.1 Dimeric Aptamers: Lane 2: FAM-Random DNA Control (2C18). Lanes 3–6: Dimeric variants of WB17/17.CD19.1_nS with varying spacer lengths (1S, 2S, 3S, 4S) exhibit clear single bands confirming their purity. Lane 7: CY3-Random DNA Control (4C18). (C) WB15/15.CD19.1 Dimeric Aptamers: Lane 2: FAM-Random DNA Control (2C18). Lanes 3–5: Dimeric variants of WB15/15.CD19.1_nS (1S, 2S, 3S) show single bands, confirming high-purity products. Lane 6: CY3-Random DNA Control (4C18). (D) WB15/17.CD19.1 Heterodimeric Aptamers: Lane 2: FAM-Random DNA Control (2C18). Lanes 3–5: Dimeric variants of WB15/17.CD19.1_nS (1S, 2S, 3S) show single sharp bands, validating successful purification. Lane 7: CY3-Random DNA Control (4C18). Dimeric aptamer concentrations: 200 nM. Gel electrophoresis ran at 120 V for 45 minutes. Gels were post-stained with SYBR Gold for 40 minutes to improve visualization. Key Insights: HPLC purification yielded high-purity dimeric aptamers, as evidenced by the absence of secondary or smeared bands. Random DNA controls from IDT exhibit expected mobility, confirming aptamer specificity and size determination. Following HPLC purification, aptamers were resolved on a 3.5% agarose gel and stained with SYBR Gold. Images were captured using a GelDoc Go Gel Imaging System (Bio-Rad).

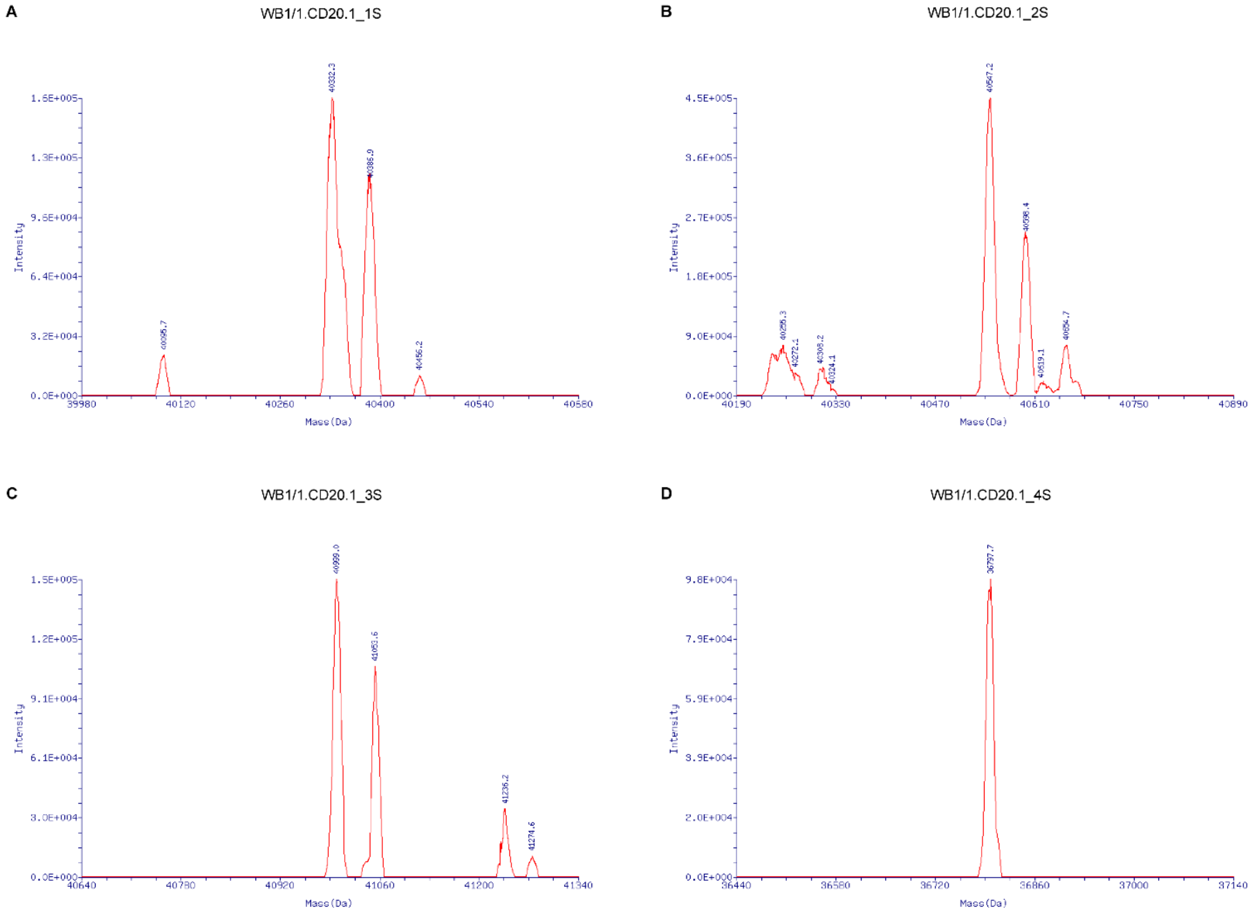


**Figure S17: Mass Spectrometry Profiles Demonstrating Dimerization Aptamer WB1/1.CD20.1_nS variants.** Mass spectrometry profiles of WB1/1.CD20.1_nS variants highlight dimerization behavior. Each spectrum corresponds to a different spacer length of homodimer WB1/1.CD20.1_nS. (A) WB1/1.CD20.1_1S dimeric peaks was observed at 40332.3 Da. (B) WB1/1.CD20.1_2S dimeric peaks were observed at 40547.2 Da. (C) WB1/1.CD20.1_3S dimeric peaks were observed at 40999.0 Da. (D) WB1/1.CD20.1_4S dimeric peaks were observed at 36797.7 Da. The x-axis represents the molecular mass (Da), and the y-axis shows the intensity of the detected ions. Peaks corresponding to dimeric forms are annotated with their respective masses. These results provide insights into the characterization of the dimeric aptamers.

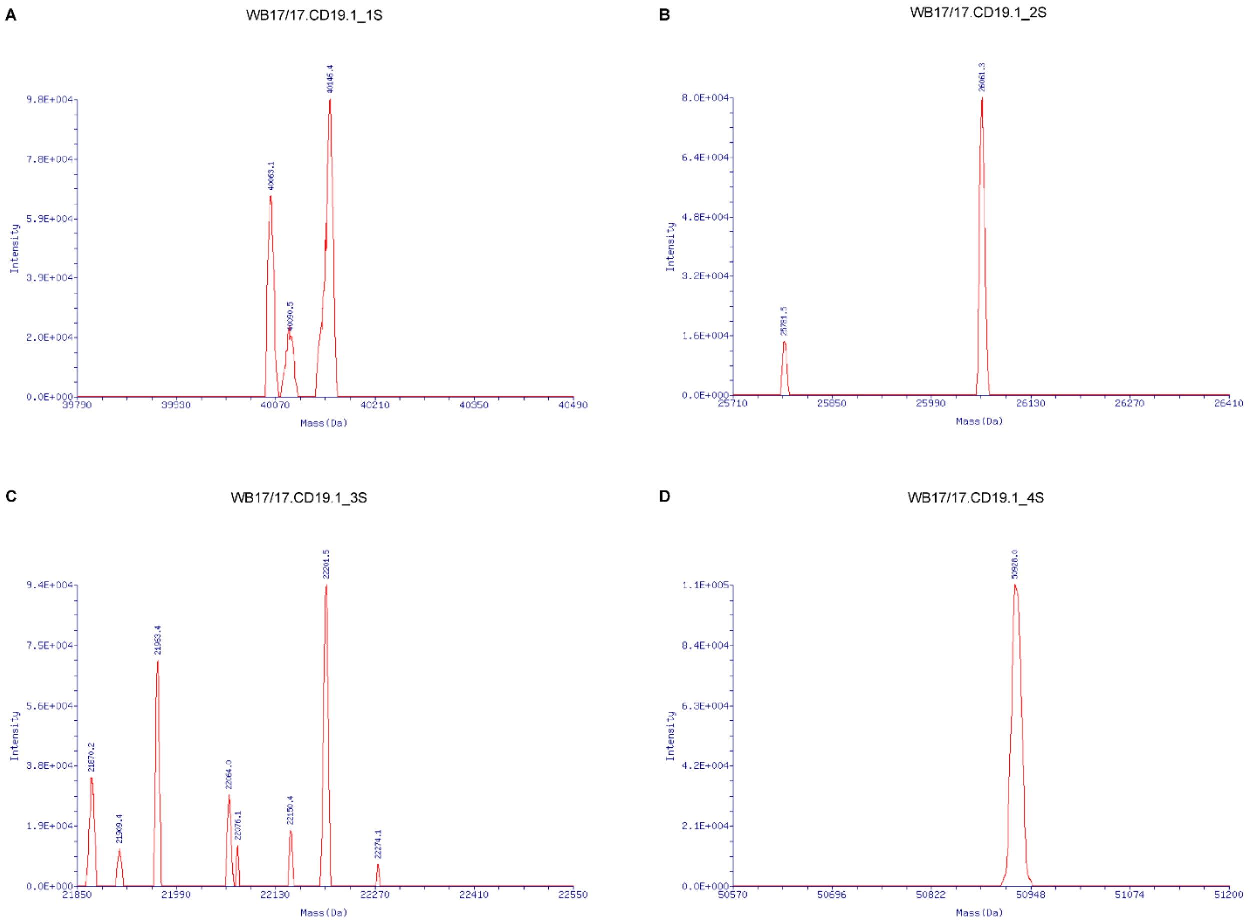


**Figure S18: Mass Spectrometry Profiles Demonstrating Dimerization Aptamer WB17/17.CD19.1_nS variants.** Mass spectrometry profiles of WB17/17.CD19.1_nS variants highlight dimerization conformation. Each spectrum corresponds to a different spacer length of homodimer WB17/17.CD19.1_nS. (A) WB17/17.CD19.1_1S dimeric peaks were observed at 40146.4 Da. (B) WB17/17.CD19.1_2S dimeric peaks were observed at 26061.3 Da. (C) WB17/17.CD19.1_3S dimeric peaks were observed at 22201.5 Da. (D) WB17/17.CD19.1_4S dimeric peaks were observed at 50928.0 Da. The x-axis represents the molecular mass (Da), and the y-axis shows the intensity of the detected ions. Peaks corresponding to dimeric forms are annotated with their respective masses. These results provide insights into the characterization of the dimeric aptamers.

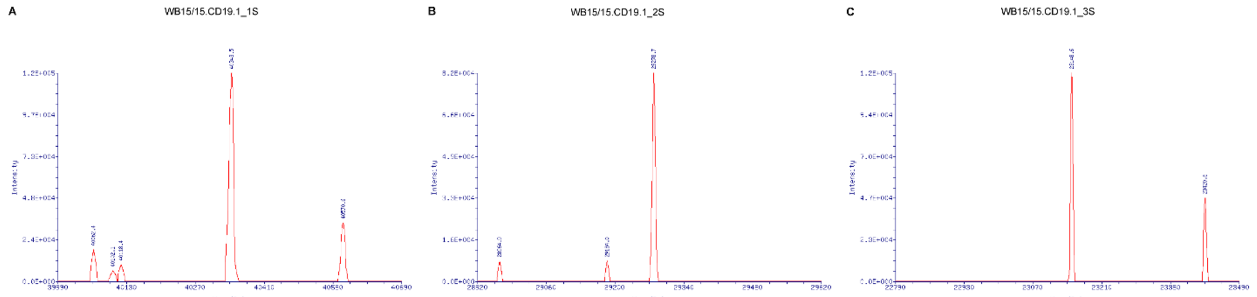


**Figure S19: Mass Spectrometry Profiles Demonstrating Dimerization Aptamer WB15/15.CD19.1_nS variants.** Mass spectrometry profiles of WB15/15.CD19.1_nS variants highlight dimerization characterization. Each spectrum corresponds to a different spacer length of homodimer WB15/15.CD19.1_nS. (A) WB15/15.CD19.1_1S dimeric peaks were observed at 40343.5 Da. (B) WB15/15.CD19.1_2S dimeric peaks were observed at 29278.7 Da. (C) WB15/15.CD19.1_3S dimeric peaks were observed at 40711.4 Da. The x-axis represents the molecular mass (Da), and the y-axis shows the intensity of the detected ions. Peaks corresponding to dimeric forms are annotated with their respective masses. These results provide insights into the characterization of the dimeric aptamers.

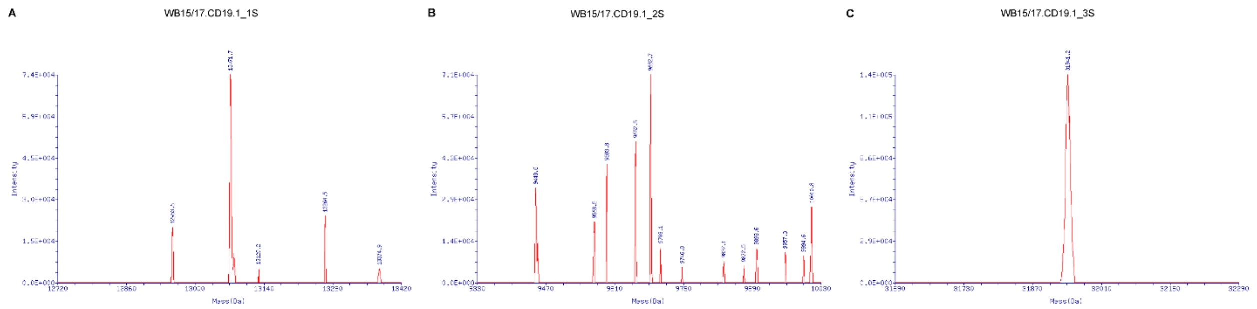


**Figure S20: Mass Spectrometry Profiles Demonstrating Dimerization Aptamer WB15/17.CD19.1_nS variants.** Mass spectrometry profiles of WB15/17.CD19.1_nS variants highlight dimerization characterization. Each spectrum corresponds to a different spacer length of homodimer WB15/17.CD19.1_nS. (A) WB15/17.CD19.1_1S dimeric peaks were observed at 13071.7 Da. (B) WB15/17.CD19.1_2S dimeric peaks were observed at 9682.7 Da. (C) WB15/17.CD19.1_3S dimeric peaks were observed at 31941.2 Da. The x-axis represents the molecular mass (Da), and the y-axis shows the intensity of the detected ions. Peaks corresponding to dimeric forms are annotated with their respective masses. These results provide insights into the characterization of the dimeric aptamers.

**Supplementary Table 1: Theoretical and Experimental Mass Data of Aptamers.** Theoretical (mass calculated) and experimental (mass found) mass data for dimeric aptamers collected using mass spectrometry. The table includes the molecule names, calculated masses (Da), and experimentally determined masses (Da). Calculated masses were derived based on the theoretical structure of each aptamer (obtained from IDT), while experimental masses were measured using mass spectrometry performed by Novatia, LLC. Aptamers were labeled with 5'-(6-FAM) or 5'-(Cy®3) fluorescent dyes are indicated in the molecule names. Discrepancies between calculated and found masses highlight potential structural modifications, dimerization, or experimental variability.

| Molecules Names | Mass Calculated [Da] | Mass Found [Da] |
|---|---|---|
| WB1/1.CD20.1_1S (FAM) | 39926.8 | 40332.3 |
| WB1/1.CD20.1_2S (FAM) | 40213.0 | 40547.2 |
| WB1/1.CD20.1_3S (CY3) | 40394.3 | 40999.0 |
| WB1/1.CD20.1_4S (CY3) | 40606.4 | 36797.7 |
| WB17/17.CD19.1_1S (FAM) | 40343.1 | 40146.4 |
| WB17/17.CD19.1_2S (FAM) | 40555.2 | 26061.3 |
| WB17/17.CD19.1_3S (CY3) | 40736.5 | 22201.5 |
| WB17/17.CD19.1_4S (CY3) | 40948.6 | 50928.0 |
| WB15/15.CD19.1_1S (FAM) | 40313.1 | 40343.5 |
| WB15/15.CD19.1_2S (FAM) | 40525.2 | 29278.7 |
| WB15/15.CD19.1_3S (CY3) | 40706.5 | 40711.4 |
| WB15/17.CD19.1_1S (FAM) | 40328.1 | 13071.7 |
| WB15/17.CD19.1_2S (FAM) | 40540.2 | 9682.7 |
| WB15/17.CD19.1_3S (FAM) | 40752.4 | 31941.2 |
